# Propidium monoazide is unreliable for quantitative live-dead molecular assays

**DOI:** 10.1101/2024.06.05.597603

**Authors:** Simerdeep Kaur, Laura Bran Ortiz, Grigorii Rudakov, Mohit S. Verma

## Abstract

Propidium monoazide (PMA) is a dye that distinguishes between live and dead cells in molecular assays like Polymerase Chain Reaction (PCR). It works by cross-linking to the DNA of cells that have compromised membranes or extracellular DNA upon photoactivation, making the DNA inaccessible for amplification. Currently, PMA is used to detect viable pathogens and alleviate systemic bias in the microbiome analysis of samples using 16S rRNA gene sequencing. In these applications, treated samples consist of different amounts of dead bacteria and a range of bacterial strains, variables that can affect the performance of PMA and lead to inconsistent findings across various research studies. To evaluate the effectiveness of PMA, we used a sensitive qPCR assay and post-treatment sample concentration to determine PMA activity accurately under varying sample conditions. We report that PMA is unreliable for viability assays when the concentration and composition of the bacterial mixture are unknown. PMA is only suitable for qualitatively assessing viability in samples containing a known number of dead microbes or extracellular DNA.

## 1. Introduction

In live pathogen detection^1–5^ and analysis of live microbial community^6–8^, researchers develop molecular viability-assays. The most commonly used method for viability assays is the treatment of samples with propidium monoazide (PMA) (Tables 1 and 2). However, researchers have used a broad range of PMA concentrations (0.5 to 100 µM, “PMA concentration selected” in Tables 1 and 2) for viable pathogen detection (Table 1) and analysis of viable microbial community (Table 2), without thorough optimization. Moreover, in an environmental or biological sample, the number of dead microbes/extracellular DNA and various strains present are crucial variables impacting PMA effectiveness^7,9–12^. Therefore, testing PMA’s response to these variables will improve our understanding of the circumstances under which this dye is effective while accounting for its limitations.

**Table 1:**
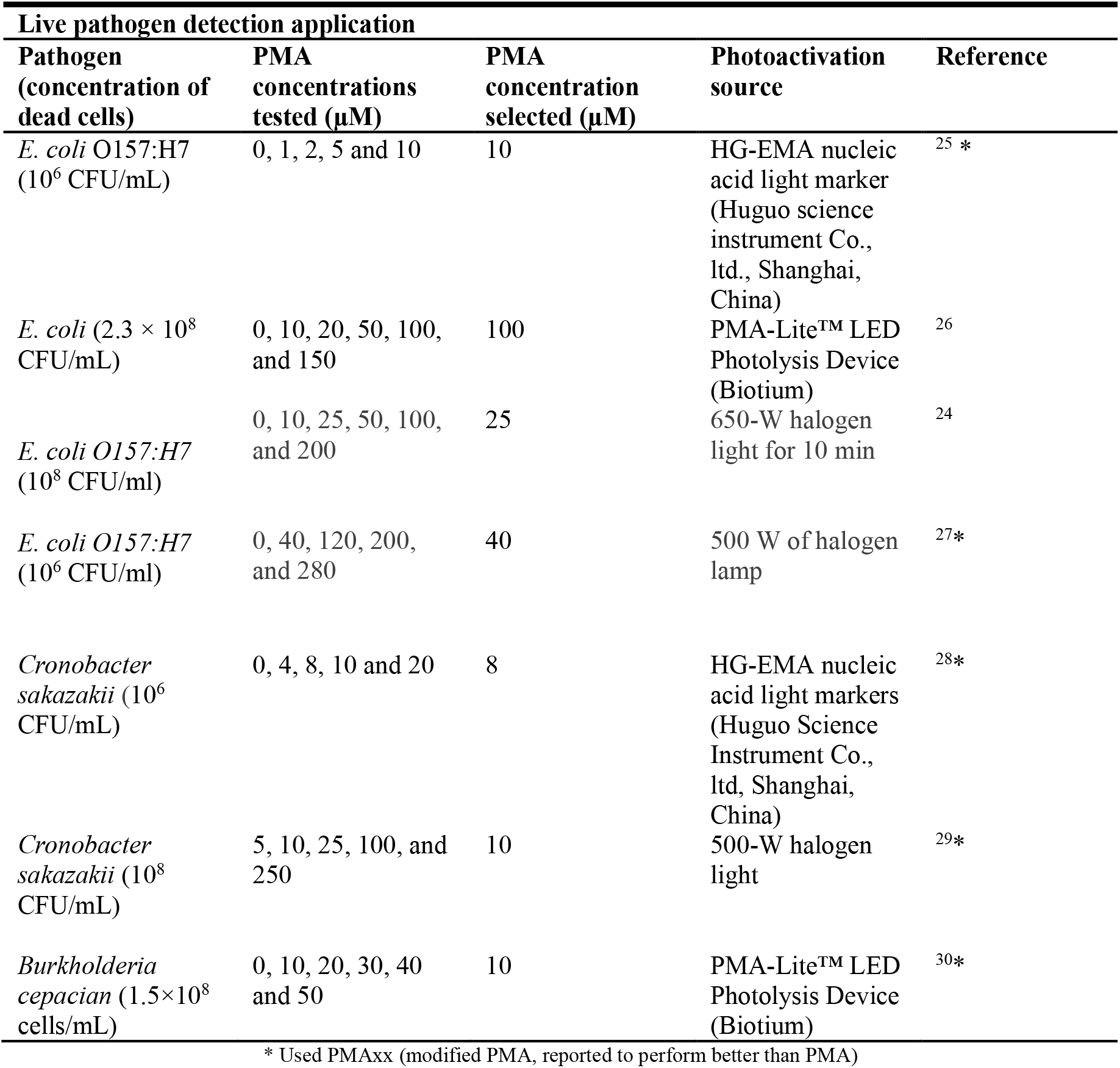
Variation in the application of propidium monoazide (PMA) as a means of detecting viable pathogens.

| **Live pathogen detection application** |  |  |  |  |
| --- | --- | --- | --- | --- |
| **Pathogen**<br>**(concentration of**<br>**dead cells)** | **PMA**<br>**concentrations**<br>**tested (μM)** | **PMA**<br>**concentration**<br>**selected (μM)** | **Photoactivation**<br>**source** | **Reference** |
| *E. coli* O157:H7<br>(10<sup>6</sup> CFU/mL) | 0, 1, 2, 5 and 10 | 10 | HG-EMA nucleic<br>acid light marker<br>(Huguo science<br>instrument Co.,<br>ltd., Shanghai,<br>China) | 25 * |
| *E. coli* (2.3 × 10<sup>8</sup><br>CFU/mL) | 0, 10, 20, 50, 100,<br>and 150 | 100 | PMA-Lite™ LED<br>Photolysis Device<br>(Biotium) | 26 |
| *E. coli* O157:H7<br>(10<sup>8</sup> CFU/ml) | 0, 10, 25, 50, 100,<br>and 200 | 25 | 650-W halogen<br>light for 10 min | 24 |
| *E. coli* O157:H7<br>(10<sup>6</sup> CFU/ml) | 0, 40, 120, 200,<br>and 280 | 40 | 500 W of halogen<br>lamp | 27* |
| *Cronobacter*<br>*sakazakii* (10<sup>6</sup><br>CFU/mL) | 0, 4, 8, 10 and 20 | 8 | HG-EMA nucleic<br>acid light markers<br>(Huguo Science<br>Instrument Co.,<br>ltd, Shanghai,<br>China) | 28* |
| *Cronobacter*<br>*sakazakii* (10<sup>8</sup><br>CFU/mL) | 5, 10, 25, 100, and<br>250 | 10 | 500-W halogen<br>light | 29* |
| *Burkholderia*<br>*cepacia* (1.5×10<sup>8</sup><br>cells/mL) | 0, 10, 20, 30, 40<br>and 50 | 10 | PMA-Lite™ LED<br>Photolysis Device<br>(Biotium) | 30* |

\* Used PMAx (modified PMA, reported to perform better than PMA)

**Table 2:**
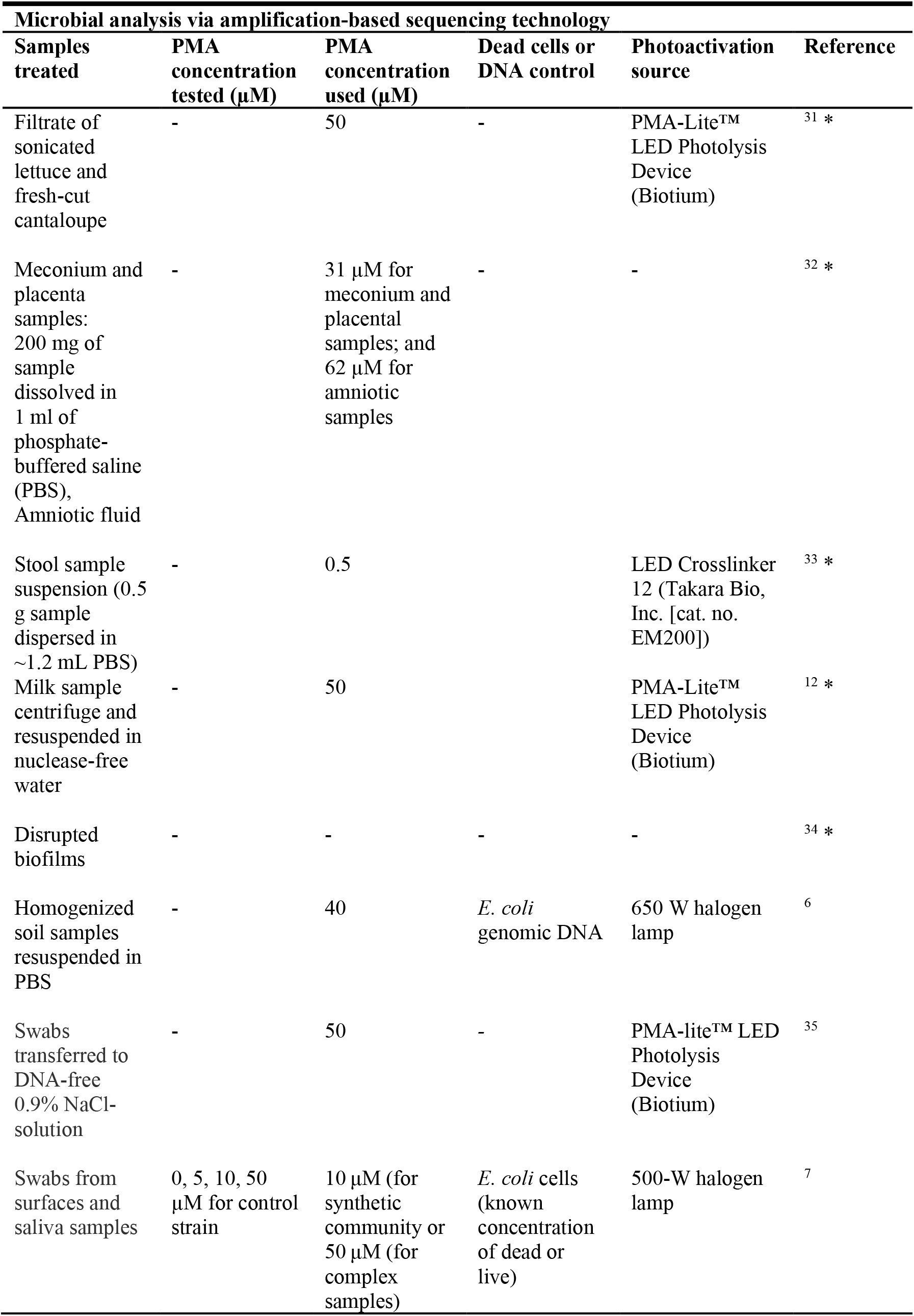
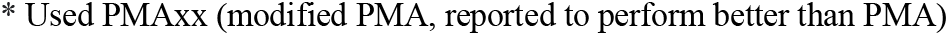
Variation in the application of propidium monoazide (PMA) as a means of viable microbiome analysis.

| **Microbial analysis via amplification-based sequencing technology** |  |  |  |  |  |
| --- | --- | --- | --- | --- | --- |
| **Samples treated** | **PMA concentration tested (μM)** | **PMA concentration used (μM)** | **Dead cells or DNA control** | **Photoactivation source** | **Reference** |
| Filtrate of sonicated lettuce and fresh-cut cantaloupe | - | 50 | - | PMA-Lite™ LED Photolysis Device (Biotium) | 31 * |
| Meconium and placenta samples: 200 mg of sample dissolved in 1 ml of phosphate-buffered saline (PBS), Amniotic fluid | - | 31 μM for meconium and placental samples; and 62 μM for amniotic samples | - | - | 32 * |
| Stool sample suspension (0.5 g sample dispersed in ~1.2 mL PBS) | - | 0.5 |  | LED Crosslinker 12 (Takara Bio, Inc. [cat. no. EM200]) | 33 * |
| Milk sample centrifuge and resuspended in nuclease-free water | - | 50 |  | PMA-Lite™ LED Photolysis Device (Biotium) | 12 * |
| Disrupted biofilms | - | - | - | - | 34 * |
| Homogenized soil samples resuspended in PBS | - | 40 | *E. coli* genomic DNA | 650 W halogen lamp | 6 |
| Swabs transferred to DNA-free 0.9% NaCl-solution | - | 50 | - | PMA-lite™ LED Photolysis Device (Biotium) | 35 |
| Swabs from surfaces and saliva samples | 0, 5, 10, 50 μM for control strain | 10 μM (for synthetic community or 50 μM (for complex samples) | *E. coli* cells (known concentration of dead or live) | 500-W halogen lamp | 7 |

\* Used PMAxx (modified PMA, reported to perform better than PMA)

**Table 3:** qPCR reaction reagents.

| Reagent (concentration) | Volume per reaction (in μL) |
| --- | --- |
| Luna® Universal Probe qPCR Master Mix (2×) | 12.5 |
| Forward primer (10μM) | 1 |
| Reverse primer (10μM) | 1 |
| Probe (10μM) | 0.5 |
| Water | 5 |
| Template (Bacterial cells or purified DNA) | 5 |
| The total volume of the reaction | 25 μL |

**Table 4:** LAMP reaction reagents.

| Reagent (concentration) | Volume per reaction (in μL) |
| --- | --- |
| NEB Warmstart Colorimetric LAMP master mix (2×) | 12.5 |
| Primer mix (10X) | 2.5 |
| Syto-9 (5mM) | 5 |
| Template (Bacterial cells or purified DNA) | 5 |
| The total volume of the reaction | 25 μL |

DNA from cells with compromised membrane integrity and extracellular DNA can survive in an environment for days to years^13^. Amplification-based detection tools for pathogens cannot distinguish between the sources (dead or live cells) of DNA and, therefore, can lead to overestimation of the pathogens or false positive results. Additionally, in DNA sequencing of microbial communities, the relic DNA contributes to systemic bias^14^. As DNA amplification and sequencing-based technologies have numerous applications^15–21^, a dependable approach for filtering out signals from dead bacteria and extracellular DNA would be valuable. One such method is PMA cross-linkage, the rampant application of PMA for identifying viable bacteria started in the early 2000s^11^. However, several studies indicate PMA treatment results in a non-specific inhibition of amplification from live cells and inefficient suppression of signal amplification from dead cells ^22–24^. For example, Wang *et al*. recently employed PMA-16S rRNA gene sequencing to measure dead cells in complex communities, and they found that PMA is useful only for qualitative analysis of these cells in such communities^7^. Furthermore, they noted that PMA treatment demonstrated a semi-quantitative capability in assessing viable cells within synthetic communities. Some studies have also reported that characteristics of nucleic acid amplification, such as amplicon size, play a role in altering PMA’s activity of inhibiting amplification signal from dead cells while retaining the signal from viable cells ^10^. Despite these reported limitations, many studies have employed this dye for viability assays (Table 1). Our study demonstrates that neglecting sample variables such as bacterial species, bacterial concentration, the sensitivity of the DNA amplification assay, and treatment parameters such as the precipitation of insoluble DNA after PMA treatment in the experimental design can result in misinterpretation of PMA activity.

In this study, we use the term “PMA cross-linkage” to represent its DNA binding aspect and “PMA activity” to denote its ability to inhibit the amplification signal from dead cells while retaining the signal from viable cells. To determine accurate PMA cross-linkage and activity with bacterial strain *E. coli* O157:H7, three strategies were employed: 1) Clear glass bottles were used for effective photoactivation, reducing day-to-day variability. 2) Following PMA treatment, a cell concentration step was carried out to eliminate the constraint of qPCR assay sensitivity with cells. 3) A DNA purification-free protocol was followed, eliminating biases from purification procedures. Upon following these startegies, we observed a “hook effect” in PMA cross-linkage with dead cells, and the optimum PMA concentration varied with the number of dead cells in the sample. This study emphasized the importance of precipitation step following cross-linkage. We also compared PMA-qPCR and PMA-LAMP assays, which provided insight into the significant role an assay’s limit of detection (LoD) plays in determining the apparent effectiveness of PMA. Furthermore, we tested a range of PMA concentrations with another bacterial species *Salmonella enterica* and found evidence of strain dependency in underestimating viable cells using PMA-qPCR.

## 2. Experimental section

### 2.1 Bacterial strains, culturing condition, and cell counting

The bacterial strain *Escherichia coli* (Migula) Castellani and Chalmers (ATCC® 35150™, serotype O157:H7) was purchased from American Type Culture Collection (ATCC). Microbiologics™ Lab-Elite™ *Salmonella enterica* subsp. enterica serovar Typhimurium ATCC™ 14028™ was purchased from Fisher Scientific (Catalog No. 23-021106). The bacteria were cultured overnight (∼16 hours) in 25 mL of Trypticase Soy Broth using a shaker-incubator at 180 RPM/37°C. 1 mL of overnight grown culture was transferred to a microcentrifuge tube and centrifuged at 10,000 RPM (using Sorvall ST4 Plus Centrifuge Series and rotor MicroClick 30 x 2 Fixed Angle Microtube Rotor) for 2 mins. The broth in the supernatant was discarded, and the pellet was either used for DNA extraction or resuspended in 1 mL Water (DNase, RNase free) for purposes like cell counting and LoD studies. For cell counting, the cells resuspended in water were diluted 10 to 100 times to bring the concentration of the cells within the working range of the bacterial cell counter: QUANTOM Tx™ microbial cell counter (Logos Biosystems, South Korea). The diluted bacteria were processed according to the manufacturer’s protocol, and the total and viable cell count was taken. The data for cell counting is shown in Figures S1, Table S1 and Figure S2, Table S2. The dilutions of cell concentrations 2×10^6^, 2×10^7^, and 2×10^8^ cells/mL were obtained for PMA treatment.

### 2.2 Genomic DNA purification and quantification

The DNA extraction was performed using the Invitrogen™ PureLink™ Genomic DNA Mini Kit129 (Fisher Scientific, K182001) following the manufacturer’s instructions. The purified DNA was quantified using Invitrogen™ Quant-iT™ dsDNA Assay Kits (Catalog number: Q33120). A 4 mL working solution was prepared by mixing 20 µL of Quant-iT™ dsDNA HS (high sensitivity) reagent in 3980 µL of Quant-iT™ dsDNA HS buffer. 100 µL of the working solution was added to 38 wells in a 96-well microplate. 5 µL of Quant-iT™ dsDNA HS standards were added to the 100 µL working solutions. The kit has 10 concentrations of dsDNA HS standards, and each concentration was run in triplicates. Three 2× serial dilutions were prepared for the purified *E. coli* O157:H7 genomic DNA and were run in the same fashion as the standards in 96 microwell plate. After sealing the plate using the microwell plate caps, it was incubated in a thermocycler at 25 °C for three cycles. After the run, the fluorescence data were plotted to get a trendline for fluorescent intensity versus the concentration of dsDNA HS standards. The equation obtained was used to estimate the concentration of the genomic DNA (Figure S3, Table S3 and S4)

### 2.3 Design and fabrication of the blue LED light device

The incubator was designed and 3D-printed in-lab with a Formlabs Form 3B 3D printer (Formlabs, Somerville, MA, USA) using Color Kit resin (Catalog # RS-F2-PKG-CR) and 0.1 mm layer thickness. The Azure color (RGB: 35, 91, 168) was achieved by following the instructions in the kit’s user manual. The electronics of the incubator include an Arduino Nano used to control the light’s color and an LED stripe (WS2812B, DigiKey) attached to the inner perimeter of the incubator (see the attached 3D model). The blue light color was achieved by programming the LED stripe to work in RGB: 0, 0, 255 regimes and powered by a 12 V power supply. The device design and sample exposure are shown in Figure S4 and S5.

### 2.4 Primer design and qPCR and LAMP reaction

LAMP primer sets (Table) for the *stxI* gene were designed using Primer Explorer v5 (http://primerexplorer.jp/lampv5e/index.html). The FASTA file of the stxI gene (NCBI accession number: M19473) was used to design the primers. Three arbitrary sets of primers (Table S5) were picked and screened for this FASTA input file. Fluorometric screening of these primers for the *stxI* gene was performed using purified genomic DNA of *E. coli* O157:H7 as the template. Three replicates of No Template Control (NTC) and test conditions were run per primer set. Test reactions had 5 µL (0.2 ng/µL) of the template, and the same volume of water was used in NTC. The genomic DNA was quantified using a Picogreen assay for dsDNA (Section 4.2). Each LAMP reaction had a final volume of 25 µL and used 2× NEB Warmstart Colorimetric LAMP master mix with 1 µM Syto-9 spiked in. Reactions were run on a qTower3G with a ramp rate of 1°C/sec. The fluorescent amplification data for primer screening is shown in Figure S6.

For qPCR assays, we used qPCR primer sets (Table S6) for the *stxI* gene published by Jinneman *et al*. in 2003, reporting high specificity and sensitivity^36^. For *Salmonella enterica*, we targeted the *invA* gene using qPCR primers and probe reported by Cheng *et al*. in 2008^37^ (Table S7).

### 2.5 Limit of detection

The limit of detection (LoD) was determined for both LAMP and qPCR assays with heat-inactivated whole cells and purified genomic DNA. The bacterial cultures were incubated overnight, followed by the transfer of the culture into a microcentrifuge tube. Subsequently, the tube was centrifuged at a speed of 10,000 RPM for a duration of 2 minutes. The supernatant was discarded, and the pellet was resuspended in water by vortexing. After proper resuspension, the culture was diluted to appropriate concentrations to determine the LoD. The calculations for the dilutions were made from the readings of the microbial cell counter for the cells. Each concentration of *E. coli* O157:H7 was vortexed for homogeneous suspension and then heated at 80 °C for 10 mins using a dry bath.

Similarly, for *Salmonella enterica*, the dilutions were heated at 70°C for 5 min. qPCR and LAMP reactions were run for the different concentrations of whole dead cell. Each concentration was run in triplicates on a 96 microwell plate. The LoD selection criterion was the lowest heat-inactivated whole cell/purified genomic DNA concentration that showed amplification in all three replicates. Similarly, the quantified genomic DNA was diluted to different concentrations after quantifying DNA using Invitrogen™ Quant-iT™ dsDNA Assay Kits (Section 4.2) and used to determine the LoD of qPCR assay.

### 2.6 PMA-qPCR and PMA-LAMP live-dead assays

1 mL of an overnight culture of *E. coli* O157:H7 or *S. enterica* was taken in a microcentrifuge tube, and the cells were pelleted down by centrifuging the tubes at 5000 RPM for 2 mins. The supernatant was discarded, and the pellet was resuspended in water. The cells were counted as mentioned in Section 4.1. A 20 mL cell suspension of desired concentration (2×10^6^, 2×10^7^, or 2×10^8^ cells/ml) was prepared, and 1 mL of this suspension was added to multiple microcentrifuge tubes (depending on the number of PMA concentrations being tested). For example, 16 1 mL cell suspension tubes were prepared to test PMA activity with 2×10^6^ cells/mL concentration. Eight tubes were for dead cells, and the other half for live cells. The dead cell tubes of *E. coli* O157:H7 were placed in the dry bath set at 80 °C/10 minutes and 70 °C/5 minutes for *S. enterica*. These are the killing conditions that provide us with dead cells (without lysing the cells). The killing of cells was confirmed by spreading plating 100 μL of heated bacteria on Trypticase Soy Agar (TSA) plates in triplicates. No growth was observed for both strains.

After heating the tubes, the samples were allowed to cool at room temperature. Eight PMA (PMAxx™ Dye, 20 mM in H2O, Biotium, Catalog No. 40069) concentrations of 0, 0.05, 0.1, 0.5, 1, 2.5, 5, and 7.5 µM were tested with both live and dead cell tubes. After adding the PMA, the tubes were incubated in the dark for 10 minutes to allow the PMA to bind to the DNA. Then 600 µL of each sample was transferred to clear glass bottles (Fisherbrand™ Class A Clear Glass Threaded Vials, Catalog No. 03-339-25B) for the photoactivation step. Our experiments provided evidence of effective photoactivation with clear glass tubes compared to natural-colored microcentrifuge tubes. The samples were exposed to blue light using the custom-made blue LED (Figure S4). After photolysis, a concentration step was performed by transferring 500 µL of the samples back into a microcentrifuge tube and centrifuging them at 10,000 RPM for 2 minutes. 250 µL of the supernatant was removed carefully without disturbing the pellet. The pellet was resuspended in the remaining 250 µL of PMA solution. The concentration step was performed to meet the LoD of the primer sets and get an accurate trend of PMA activity. The samples were used for qPCR and LAMP as described in Section 4.4.

## 3. Results and Discussion

### 3.1 PMA cross-linkage shows “hook effect” like activity trend

In the preliminary trials, we screened PMA concentrations of 5 µM or higher based on reported effectiveness in the literature (Tables 1 and 2). We observed inhibition of amplification at these reported concentrations. However, after introducing a step to concentrate the samples post-PMA treatment through centrifugation and resuspending them in half of their original volume, the qPCR assays detected amplification in dead *E. coli* O157:H7 at PMA concentration 5 µM or higher (Figure 1A). Therefore, for samples that were not concentrated, the assay’s limited sensitivity compensated for the suboptimal performance of PMA, owing to the template concentration falling below the qPCR assay’s limit of detection (LoD), which is established at 200 dead cells per reaction or 200 dead cells per 5 µL of sample volume, as demonstrated in Figure S7.

**Figure 1:**
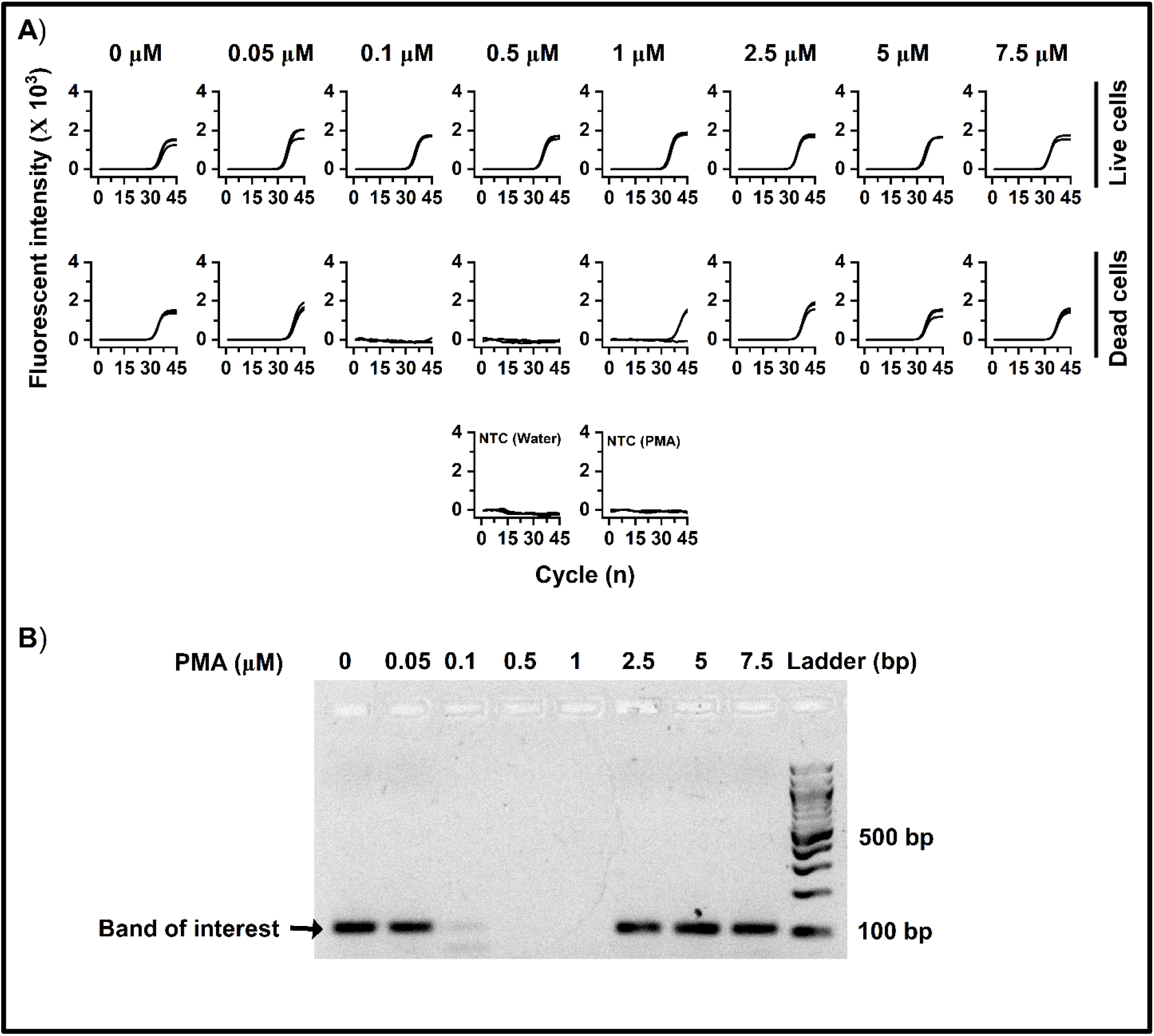
PMA-qPCR results for 2×10^6^ cells/mL treated at different concentrations of PMA A) The plots in the first row show the fluorescent qPCR amplification data for live cells treated with varying concentrations of PMA (labeled at the top). The graph in the second row represents the same data for dead cells. The plots in the third row are for controls, with water and PMA solution replacing the samples in the reaction. Each sample was tested in triplicates. B) The gel electrophoresis results display one of the replicates of PMA-qPCR reactions for dead cells. Each well is labeled with the corresponding PMA concentration used during treatment. NTC: No template control, bp: base pair

With concentrated samples, we explored the efficacy of PMA across a spectrum of concentrations from 0.05 to 1 µM. The results, illustrated in Figure 1A, revealed a discernible elevation in the Ct values correlating with increased concentrations of PMA for non-viable cells. Specifically, the Ct value increased from 38.673 at a PMA concentration of 0.05 µM to a scenario of non-amplification, which we infer as an assumed Ct value of 45, upon treatment with 0.5 µM PMA. Through this experimental approach, we identified that the effective concentration window of PMA, where dead cells exhibited no amplification, spanned from 0.1 µM to 1 µM. However, surpassing this concentration threshold led to significant amplification signals (Ct value of 36.87 at 2.5 µM PMA) from dead cells, suggesting a pattern akin to the “hook effect”, a well-documented phenomenon in bioanalytical assays^38^.

We validated the amplification of PMA-treated dead cells using gel electrophoresis, analyzing one of the three qPCR assay replicates. Dead cells treated with 2.5 to 7.5 µM PMA had amplicon bands comparable to the non-treated dead cells (no PMA), confirming the “hook effect” of PMA (Figure 1B). At lower concentrations of the template (Figure 1A and B, 0.1 to 1 µM PMA, dead cells), inconsistency among qPCR replicates is commonly reported^39^. Live cells showed consistent amplification among PMA-treated and non-treated samples (Figure 1A). We removed free PMA after photoactivation by centrifuging the samples for this experiment. To ensure reproducibility, we repeated this experiment three times.

DNA intercalating dyes are known to induce structural modifications in DNA due to their intercalation mechanism, which requires DNA unwinding to accommodate the dye between base pairs^40,41^. Therefore, at higher PMA concentrations, it is possible that decrease in Ct value is because of excessive intercalation that causes localized distortions in the DNA structure, making it accessible for amplification. These distortions can disrupt hydrogen bonding interactions and potentially increase the accessibility of DNA for amplification processes.

### 3.2 PMA cross-linkage varies with different concentrations of dead cells

Having observed the optimal PMA concentration range of 0.5 to 1 µM (Figure 1A) with 2×10^6^ cells/mL, our next objective was to determine whether this range suits other concentrations of dead *E. coli* O157:H7 cells.To achieve this, we evaluated different PMA concentrations across a spectrum of live and dead cell densities. For the experiments under this section, we opted not to remove free PMA after photoactivation because we observed that the removal of free PMA did not impact the qPCR assay ^42^. We heated the samples at 90°C for 5 minutes to extract the DNA from live and dead cells and use them as a template for qPCR.

With a cell concentration of 2×10^6^ cells/mL (Figure 2A), we observed the highest cycle threshold (Ct) value with dead cell samples at 0.5 µM PMA. The Ct value decreased with increased PMA concentration beyond 0.5 µM PMA. We observed a similar activity trend with a cell concentration of 2×10^7^ cells/mL (Figure 2B), but the optimum PMA concentration shifted to 2.5 µM. On testing cell concentration 2×10^8^ cells/mL (Figure 2C), the highest Ct value with dead cells moved to 10 µM PMA. Moreover, the highest Ct value at optimum PMA concentration also decreased with increasing cell concentration. The corresponding fluorescent qPCR data for Figure 2 is in Figures S8, S9, and S10.

**Figure 2:**
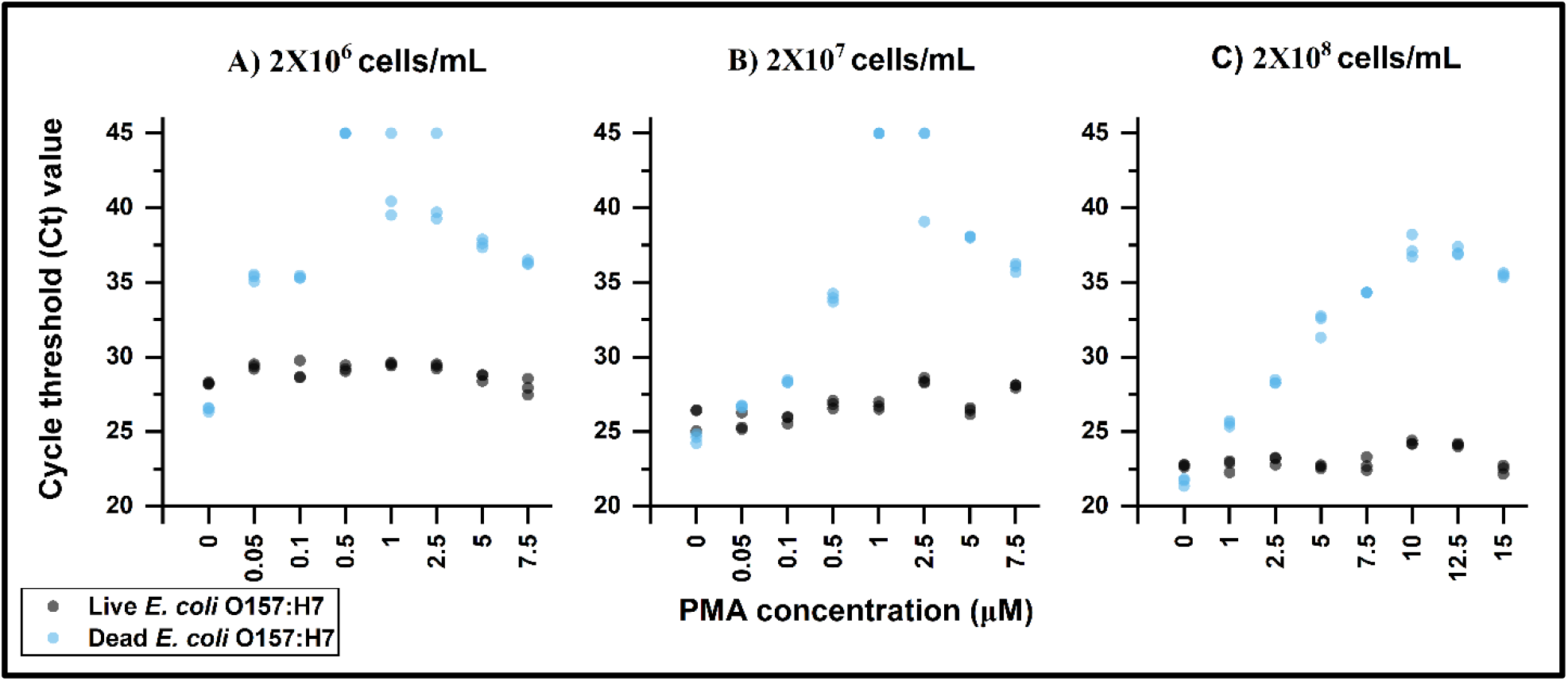
qPCR analysis of PMA treatment efficacy across varied *E. coli* O157:H7 cell concentrations: The plots show the cycle threshold (Ct) values obtained from qPCR analysis of PMA-treated cells. Each sample was tested in triplicate. The x-axis represents the range of PMA concentrations used, while the y-axis represents the Ct values. Three different cell concentrations were tested: A) 2×0^6^ cells/mL, B) 2×10^7^ cells/mL, and C) 2×10^8^ cells/mL. In cases where no amplification was observed, the Ct value was assigned as the last cycle (45).

Our observations might explain other published studies, e.g., in the study by Stinson *et al*., a PMA concentration of 50 µM was employed to assess the viable milk microbiome^12^. The findings demonstrated a notable reduction in DNA yield from the samples treated with PMA. However, it is important to note that they did not spike control dead bacteria or extracellular DNA in the test samples. As a result, it cannot conclusively determine the extent of PMA treatment.Wang *et al*. reported the ineffectiveness of PMA treatment in removing the signal from control bacteria in complex samples^7^. Furthermore, when attempting to remove relic DNA from soil samples, Carini *et al*. observed that samples with high relic DNA concentration showed only a reduced signal from spiked extracellular DNA (control)^6^. In both these studies, PMA underperformed when the dead cell or relic DNA number changed significantly.

Nocker *et al*. observed that the binding and photo-induced cross-linkage of PMA rendered the DNA insoluble, leading to its loss along with cellular debris during the DNA purification procedure^11^. In our experiments (Figure 2), we extracted DNA by heat lysis before introducing the samples to qPCR reactions. However, we deliberately excluded DNA purification steps from our analysis. This omission allowed us to perform post-treatment cell concentration, eliminate biases from purification protocols, and facilitate the investigation of PMA cross-linkage.

Additionally, researchers have reported PMA underperformance, even after DNA extraction and purification protocols^6,7^. The PMAxx protocol from Biotium specifies that insoluble DNA is removed by precipitation during the DNA purification process.We tested precipitation by centrifugating (10,000 RPM for 5 minutes) the samples to remove insoluble fractions from PMA-treated live and dead cells (2×10^8^ cells/mL) after heat lysis. We observed a complete elimination in PCR signal with the dead cell samples beyond 7.5 µM PMA, making 10 µM PMA the optimum PMA concentration (Figure S11). However, this precipitation step also led to the loss of DNA from live cells (23.3 Ct value with no PMA and 29.4 Ct value with 10 µM PMA), indicating that the removal of the signal is not specific to dead cells, making PMA unreliable for quantitative analysis of viability. These findings also suggest that the precipitation step plays a crucial role in the effectiveness of PMA and should not be solely reliant on DNA purification steps. Therefore, a separate optimization of the precipitation step is essential.

### 3.3 PMA cross-linkage shows “hook effect” with purified genomic DNA

PMA-qPCR of the cell samples comes with some skepticism, mainly because the turbidity of the bacterial cell suspension can impact the blue light exposure^43^. Furthermore, as we did not purify DNA, the cellular biomass becomes a significant biological variable across different cell concentrations in Section 2.2. Therefore, we can potentially attribute the observed variation in Ct values to qPCR variability influenced by biomass. To test this factor, we treated purified genomic *E. coli* O157:H7 DNA to test these factors with PMA. When analyzing genomic DNA, we noticed a similar phenomenon to the “hook effect” observed with cells. Specifically, an optimal PMA concentration resulted in decreased Ct values, followed by a decline in Ct values beyond this optimal concentration (Figures 3A and B). On comparing DNA concentrations with a 10 times difference, we also observed a shift in optimum PMA concentration from 1 µM to 2.5 µM. The LoD of the qPCR assay with genomic DNA is 1 copy/reaction or 1 copy/5 μL of sample volume (Figure S12). In addition to exhibiting the “hook effect”, PMA could not remove the amplifications at any concentration.

**Figure 3:**
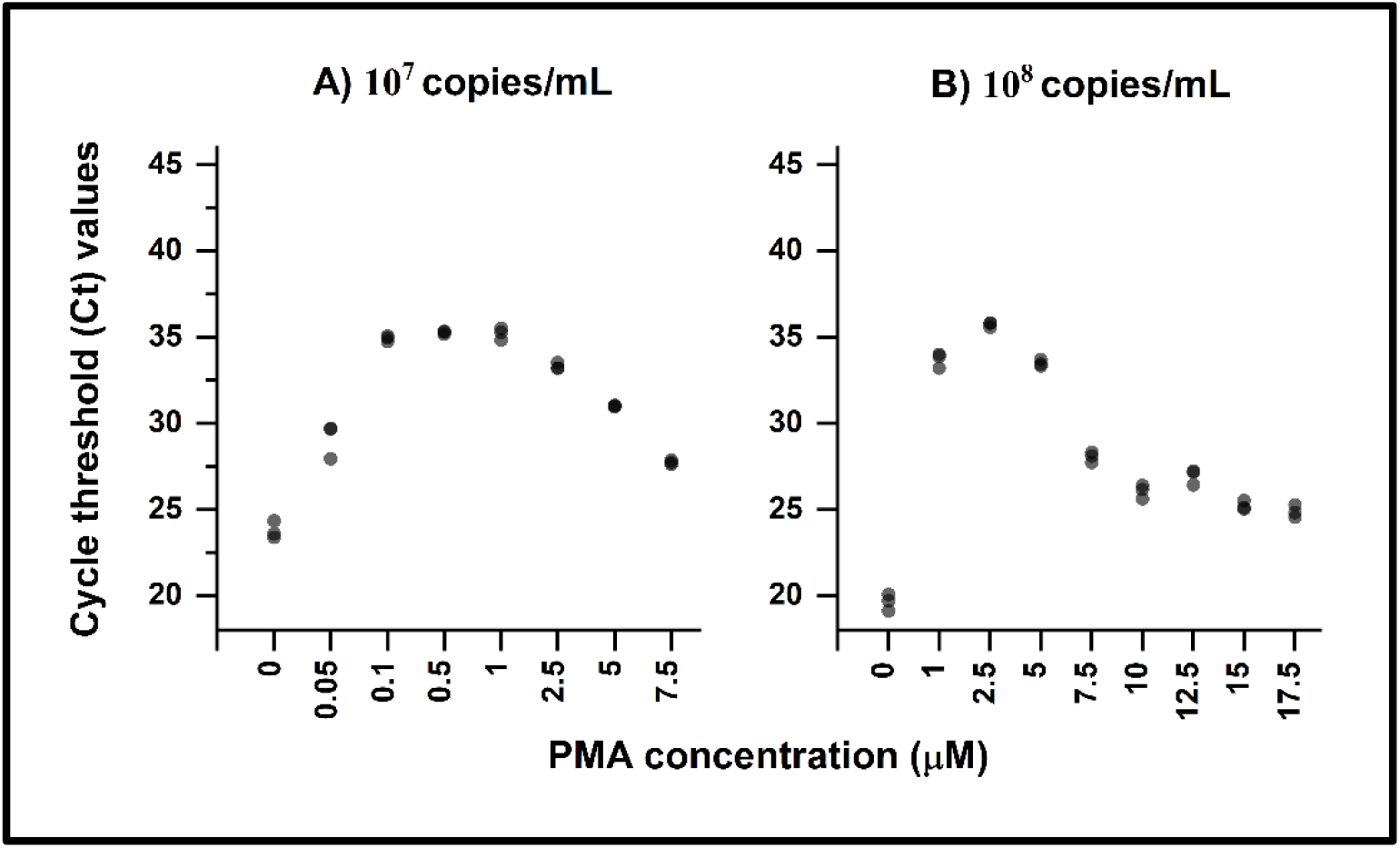
qPCR analysis of PMA treatment efficacy across different genomic DNA concentrations of *E. coli* O157:H7: The plots display the cycle threshold (Ct) values obtained through qPCR analysis of PMA-treated genomic DNA. Each sample was tested in triplicate. The x-axis represents the range of PMA concentrations applied, while the y-axis represents the Ct values. Two DNA concentrations were examined: A) 10^7^ copies/mL and B) 10^8^ copies/mL.

### 3.4 The custom blue LED has similar photoactivation as PMA-lite™ LED Photolysis Device (Biotium)

After eliminating the possibility of turbidity of cells affecting photoactivation, we wanted to test if PMA activity depends on the blue light source. Since all our studies were performed with our custom-built blue LED device, comparing PMA activity with a commercial device was important. We observed a similar PMA activity trend with PMA-Lite™ LED Photolysis Device (Biotium) (the commercial device designed for photoactivation) and our custom blue LED (Figure 4). The Ct values vary among the light exposure treatments in Figure 4, but the activity trend of PMA remains the same.

**Figure 4:**
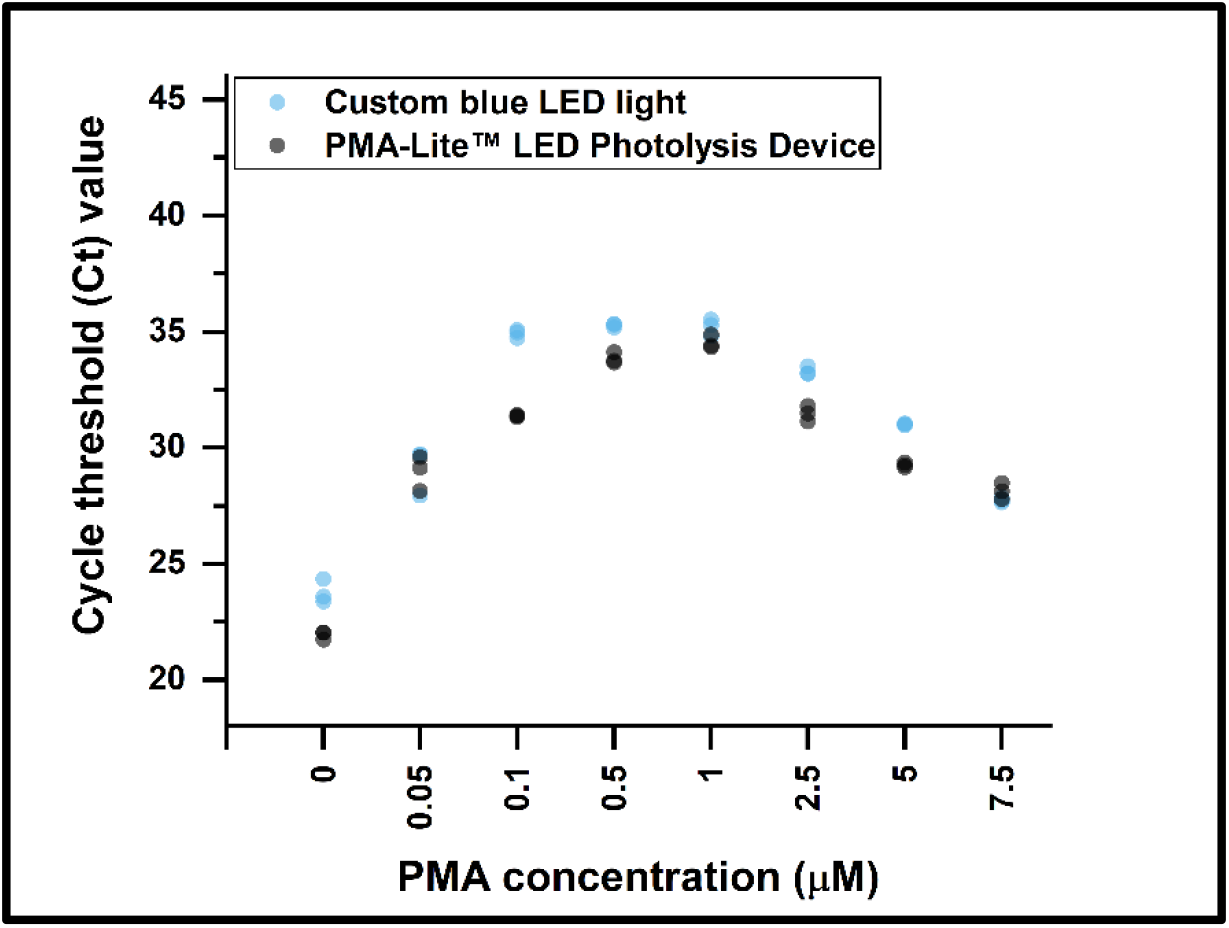
qPCR analysis of PMA treatment efficacy with different sources of blue light: Cycle threshold (Ct) (y-axis) values generated with qPCR for PMA-treated genomic DNA (10^7^ copies/mL); each sample was run in triplicates

### 3.5 Sensitivity of the DNA amplification assay plays a role in determining PMA activity

Isothermal amplification assays, such as loop-mediated isothermal amplification (LAMP), with PMA, offer a prospect for developing field-deployable live pathogen detection assays^2^. We compared the qPCR results for PMA-treated samples with an isothermal DNA amplification method. With LAMP assays, we observed a gradual increase in the time threshold (Tt) value, followed by no amplification, upon increasing concentrations of PMA in the assays. Notably, PMA-treated dead cells at 2×10^6^ cells/mL exhibited no amplification even at a low PMA concentration of 0.1 μM (Figure S13). Similarly, dead cells at 2×10^8^ cells/mL showed no amplification beyond a PMA concentration of 5 μM (Figure S14), observing no hook effect. We can attribute the disparity in PMA activity between qPCR and LAMP assays to the significant difference in the limit of detection (LoD) of the assays. The LoD for LAMP assays using dead cells as templates is 1000 dead cells/reaction or 1000 dead cells/5 μL of sample volume (Figure S15), whereas for qPCR, it is 200 dead cells/reaction or 200 dead cells/5 μL of sample volume (Figure S7). Importantly, the decreased sensitivity of the LAMP assay renders the “hook effect” invisible. Therefore, carefully considering the assay’s LoD is essential when assessing PMA activity to avoid potential misinterpretation of results.

Studies have reported a gradual increase in Tt value or Ct value as a factor of PMA concentration^25,28^. In their research, Rani *et al*. noticed a variation in the optimum PMA concentration when utilizing PMA-treated samples as the template for DNA amplification assays with two distinct primer sets^26^. Numerous studies have also indicated that the amplicon size can influence the inconsistent results obtained from PMA treatment^10,29,44,45^. Therefore, the variables associated with PMA treatment, sample variables, and the parameters of the amplification reactions control the effectiveness of PMA activity.

### 3.6 Underestimation of live cells depends on the bacterial strain

In our implementation of PMA treatment combined with cell concentration to *S. enterica*, we observed that the optimal range for PMA concentration is 1 to 2.5 µM (Figure 5). Similar to the “hook effect” in *E. coli* (Figure 1), we observed the presence of this phenomenon in *S. enterica*, as indicated by the drop in Ct values at concentrations of PMA exceeding 7.5 µM (Figure 5). Interestingly, unlike the outcomes observed in *E. coli*, PMA consistently underestimated the presence of live cells in *S. enterica* at all tested concentrations. For instance, the Ct value increased from 23.3 for untreated live cells to 30.3 with 0.5 µM PMA-treated live cells. Therefore, we can conclude that PMA exhibits a strain-specific tendency to underestimate the presence of live cells. The LoD of the qPCR assay with heated whole dead *S. enterica* is 250 cells/reaction or 250 cells/5 µL of the sample volume (Figure S16). The corresponding fluorescent qPCR data for Figure 5 is in Figure S17. It is possible to prevent the underestimation of live cells by limiting and optimizing the PMA binding and blue light exposure time. However, in the case of PMA-16S rRNA technology, the uniform treatment of bacteria within a diverse community may lead to an underestimation of viable cells for certain bacteria species. The variability with live cells among strains is a critical factor that needs to be considered during microbial analysis using PMA-16s rRNA gene sequencing.

**Figure 5:**
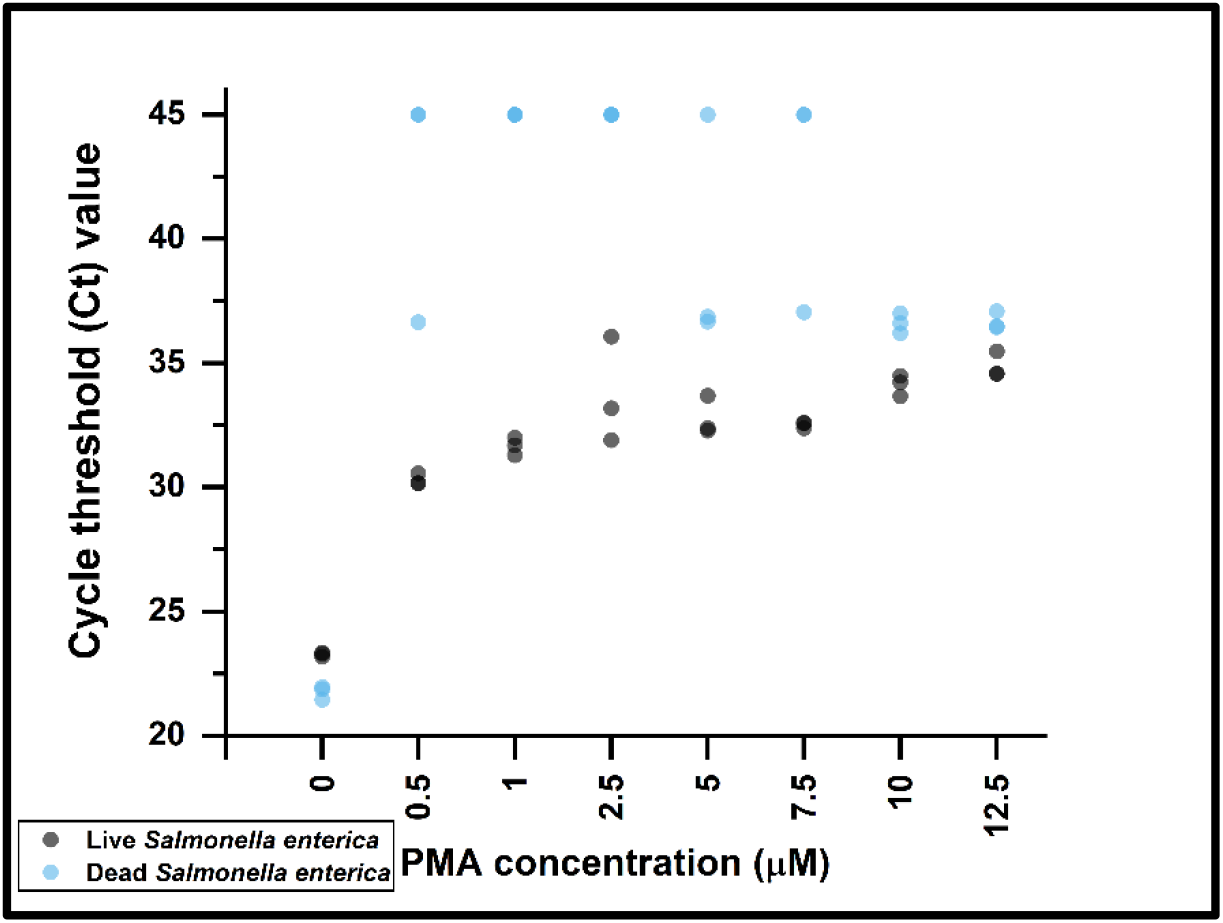
PMA-qPCR results for *Salmonella enterica* treated at different PMA concentrations: Cycle threshold (Ct) (x-axis) value for cell concentration 2×10^6^ cells/mL at PMA concentrations (y-axis). For samples showing no amplification, the Ct value was assumed to be the last cycle (45)

## 4. Conclusions

Three key features helped us achieve the unique trends in PMA cross-linkage: 1) We gave considerable attention to factors related to sample variability, PMA treatment, and amplification assays. By meticulously considering these factors, we ensured a robust experimental setup, minimizing potential biases and inaccuracies in our findings. 2) Post-treatment with PMA, we concentrated the samples to preclude any possible misconceptions concerning PMA efficacy attributable to the amplification assay’s insufficient sensitivity. 3) We distinguished PMA cross-linkage from precipitation, a critical aspect that provided valuable insights into the frequent underperformance of PMA reported in numerous studies. This enabled us to develop a potential explanation for the observed phenomena, shedding light on an area subject to ambiguity and confusion.

However, it is important to acknowledge two limitations of our study: 1) We did not compare our procedure of PMA treatment in this study to the one reported in the literature (PMA treatment followed by DNA purification with commercial kits). 2) While our claims related to 16s rRNA gene sequencing provide valuable insights, it is essential to recognize that the steps involved in this technology differ from DNA amplification assays. Therefore, additional validation is necessary to solidify the reliability and accuracy of our results to 16s rRNA gene sequencing.

Overall, our study has significantly contributed to clarifying the ambiguities surrounding the usability of PMA. The insights gained from our research highlight essential additional steps researchers should consider when optimizing the assay for various applications. We recognize that the challenges associated with PMA usage make it unreliable for quantitative analysis.

## 5. Supporting information

The Supporting Information is available with this publication.

## 6. Data availability statement

The data sets generated and/or analyzed during the current study are available in the Mendeley data repository at DOI: 10.17632/sp8k2p9nx6.1

## 7. Acknowledgments

This research was supported by the Center for Food Safety Engineering at Purdue University, funded by the U.S. Department of Agriculture, Agricultural Research Service, under Agreement No. 59-8072-1-002. Any opinions, findings, conclusion, or recommendations expressed in this publication are those of the author(s) and do not necessarily reflect the view of the U.S. Department of Agriculture. The work was also partially supported by the Center for Produce Safety (CPS Award Number: 2021CPS12), the California Department of Food and Agriculture (CDFA Agreement No. 20-0001-054-SF), and the U.S. Department of Agriculture’s (USDA) Agricultural Marketing Service (USDA Cooperative Agreement No. USDA-AMS-TM-SCBGP-G-20-0003). Any opinions, findings, conclusions, or recommendations expressed in this publication or audiovisual are those of the author(s) and do not necessarily reflect the views of The Center for Produce Safety, the California Department of Food and Agriculture, or the U.S. Department of Agriculture’s (USDA) Agricultural Marketing Service. The project entitled Field evaluation of microfluidic paper-based analytical devices for microbial source tracking was funded in whole or in part through a subrecipient grant awarded The Center for Produce Safety through the California Department of Food and Agriculture 2020 Specialty Crop Block Grant Program and the U.S. Department of Agriculture’s (USDA) Agricultural Marketing Service. We are grateul to Jiangshan Wang for helping in improving the writing quality of this manuscript.

## 8. Declaration of AI and AI-assisted technologies in the writing process’

During the preparation of this work the authors used Grammarly (https://grammarly.com/) and ChatGPT (https://chat.openai.com/) in order to check for grammar errors and improve the academic writing language. After using this tool/service, the authors reviewed and edited the content as needed and take full responsibility for the content of the publication.

## 9. Conflict of Interest

M.S.V. has an interest in Krishi Inc., which is a startup that is interested in commercializing technologies developed here. This work was not funded by Krishi Inc.

