## Supporting Information for "Propidium monoazide is unreliable for quantitative live-dead molecular assays"

For

1. **Supporting data and information for the material and methods section**

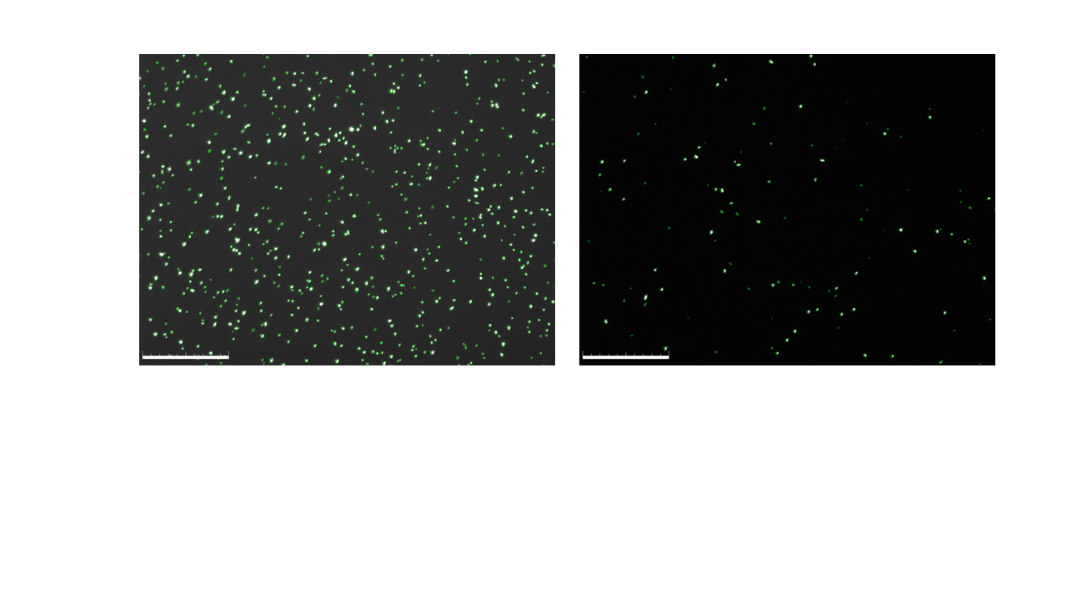

Figure S1: Microbial cell counter image depicting the stained culture of *E. coli* O157:H7, diluted 10 times. The left image illustrates staining for the total cell count, while the right image depicts staining for the count of viable cells. A scale bar of 100 µm is provided for reference.

Table S1: Microbial cell counter readings acquired for *E. coli* O157:H7, as depicted in Figure S1.

| **Total cell count** | |
| --- | --- |
| Replicate 1 | 2.67 x 10e8 cells/mL |
| Replicate 2 | 2.38 x 10e8 cells/mL |
| **Live cell count** | |
| Replicate 1 | 3.24 x 10e7 cells/mL |
| Replicate 2 | 3.51 x 10e7 cells/mL |

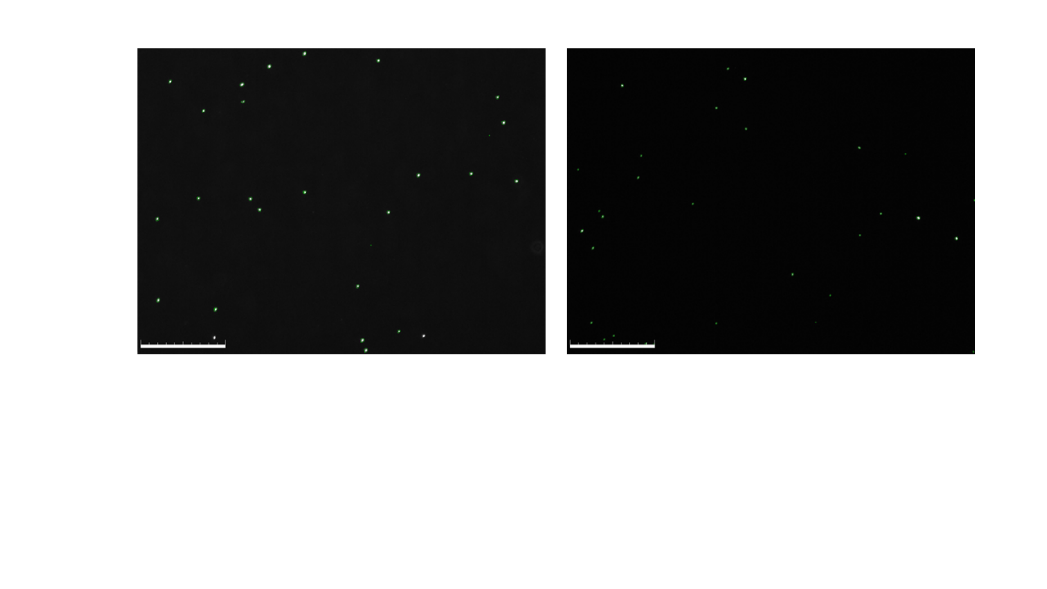

Figure S2: Image of microbial cell counter data illustrating the culture of *Salmonella enterica*, diluted 100 times. The left image presents staining for the total cell count, while the right image showcases staining for the count of viable cells. A scale bar of 100 µm is included for scale reference.

Table S2: Microbial cell counter readings acquired for *Salmonella enterica*, as depicted in Figure S2.

| **Total cell count** | |
| --- | --- |
| Replicate 1 | 4.00 x 10e6 cells/mL |
| Replicate 2 | 4.38 x 10e6 cells/mL |
| **Live cell count** | |
| Replicate 1 | 4.30 x 10e6 cells/mL |
| Replicate 2 | 6.15 x 10e6 cells/mL |

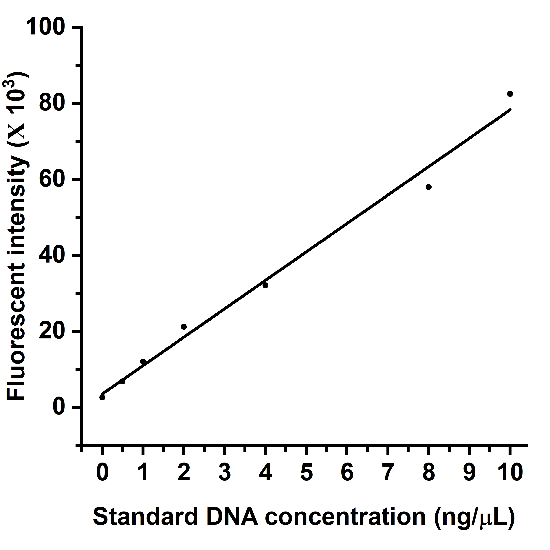

Figure S3: A) Standard curve plot for quantifying purified genomic DNA of *E. coli* O157:H7 using Invitrogen™ Quant-iT™ dsDNA Assay.

Table S3: The equation obtained for calculating DNA concentration in the sample derived from the calibration curve in Figure S3

| Equation | y = a + b*x |
| --- | --- |
| Intercept | 3543.01116 ± 1810.3011 |
| Slope | 7482.06024 ± 351.90118 |
| Residual Sum of Squares | 5.7185E7 |
| Pearson's r | 0.99452 |
| R-Square (COD) | 0.98906 |
| Adj. R-Square | 0.98687 |

Table S4: Fluorescent intensities of three dilutions of *E. coli* O157:H7 genomic DNA in Invitrogen™ Quant-iT™ dsDNA Assay. The DNA sample concentration was calculated by plugging in the equation y=7482.06x+3543. The original concentration was calculated by multiplying the sample concentration by the dilution factor. The average concentration of genomic DNA calculated= 68.99278 (ng/µL)

|  | Replicate 1 | Replicate 2 | Replicate 3 | Average | Sample concentration (ng/µL) | Original concentration (ng/µL) |
| --- | --- | --- | --- | --- | --- | --- |
| Dilution 1 | 57091.23 | 55873.89 | 53913.95 | 55626.36 | 6.961061 | 69.61061 |
| Dilution 2 | 30766.27 | 30191.37 | 26586.16 | 29181.27 | 3.426614 | 68.53227 |
| Dilution 3 | 16857.77 | 17515.19 | 14883.57 | 16418.84 | 1.720886 | 68.83545 |

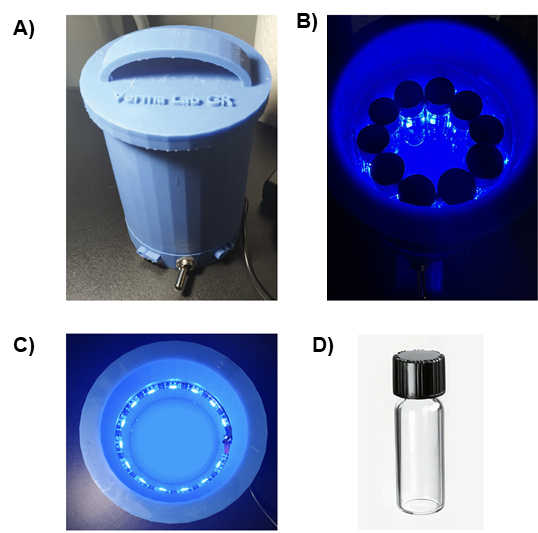

Figure S4: Photoactivation setup: A) Custom-built LED light incubator outer structure. B) Bacterial samples in clear glass vials in a custom-built LED device exposed for 15 minutes in the dark. C) Top view of the custom-built blue LED device without the lid. D) Clear glass vials used for photoactivation (Fisherbrand™ Class A Clear Glass Threaded Vials).

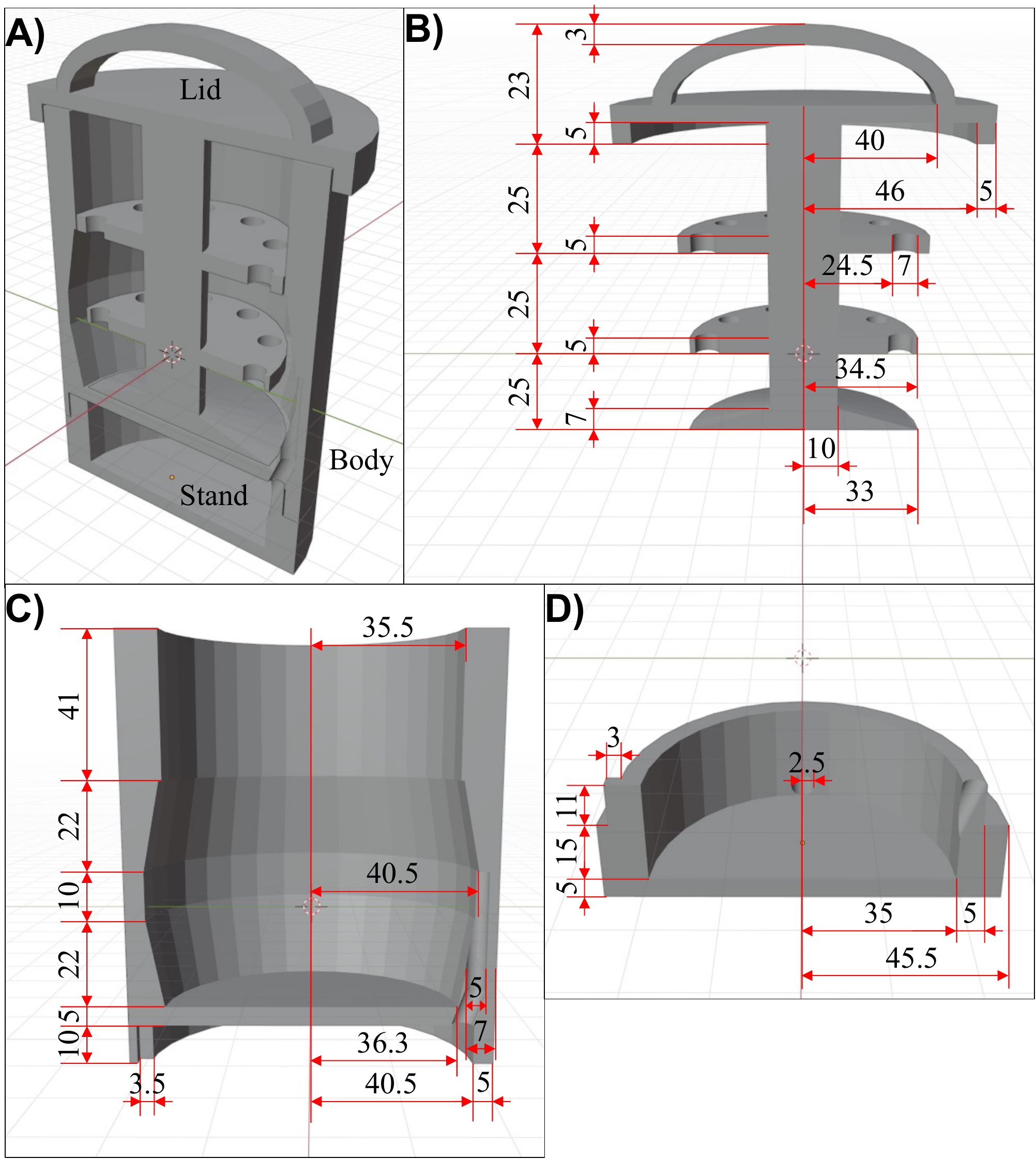

Figure S5: Vertical cross-section CAD (computer-aided design) models of the custom-built LED light: A) Raw figure of the incubator body, tube holder, and base in assembled state. B) Tube holder dimensions C) Body dimensions D) Base dimensions of the incubator body.

Table S5: Sequence of primers screened for LAMP assay for *stx1* gene of *E. coli* O157:H7

| **Oligo sequence (5'- 3')** | **Oligo name** |
| --- | --- |
| TGATTTTTCACATGTTACCTTTC | EC.stx1.1_F3 |
| TAACATCGCTCTTGCCAC | EC.stx1.1_B3 |
| CCTGCAACACGCTGTAACGTCAGGTACAACAGCGGTTA | EC.stx1.1_FIP |
| AGTCGTACGGGGATGCAGATAGTGAGGTTCCACTATGC | EC.stx1.1_BIP |
| GTATAGCTACTGTCACCAGACAATG | EC.stx1.1_LF |
| AAATCGCCATTCGTTGACTACT | EC.stx1.1_LB |
| AGCGTGGCATTAATACTGAA | EC.stx1.2_F3 |
| TCATTTTACCCCCTCAACT | EC.stx1.2_B3 |
| CCGGACACATAGAAGGAAACTCTCATCATCATGCATCGCG | EC.stx1.2_FIP |
| GAGTCCGTGGGATTACGCACGCTAATAGTTCTGCGCATCA | EC.stx1.2_BIP |
| ATGCCATTCTGGCAACT | EC.stx1.2_LF |
| AAAATATTGTGGGATTCATCCACTC | EC.stx1.2_LB |
| TTACCTTTCCAGGTACAACA | EC.stx1.3_F3 |
| GTAAAGCTTCAGCTGTCAC | EC.stx1.3_B3 |
| TACGACTGATCCCTGCAACACGGTTACATTGTCTGGTGAC | EC.stx1.3_FIP |
| TGGATTTAATGTCGCATAGTGGAACAAACCGTAACATCGCTCTTG | EC.stx1.3_BIP |
| CGCTGTAACGTGGTATAGCTACT | EC.stx1.3_LF |
| CTCACTGACGCAGTCTGTGG | EC.stx1.3_LB |

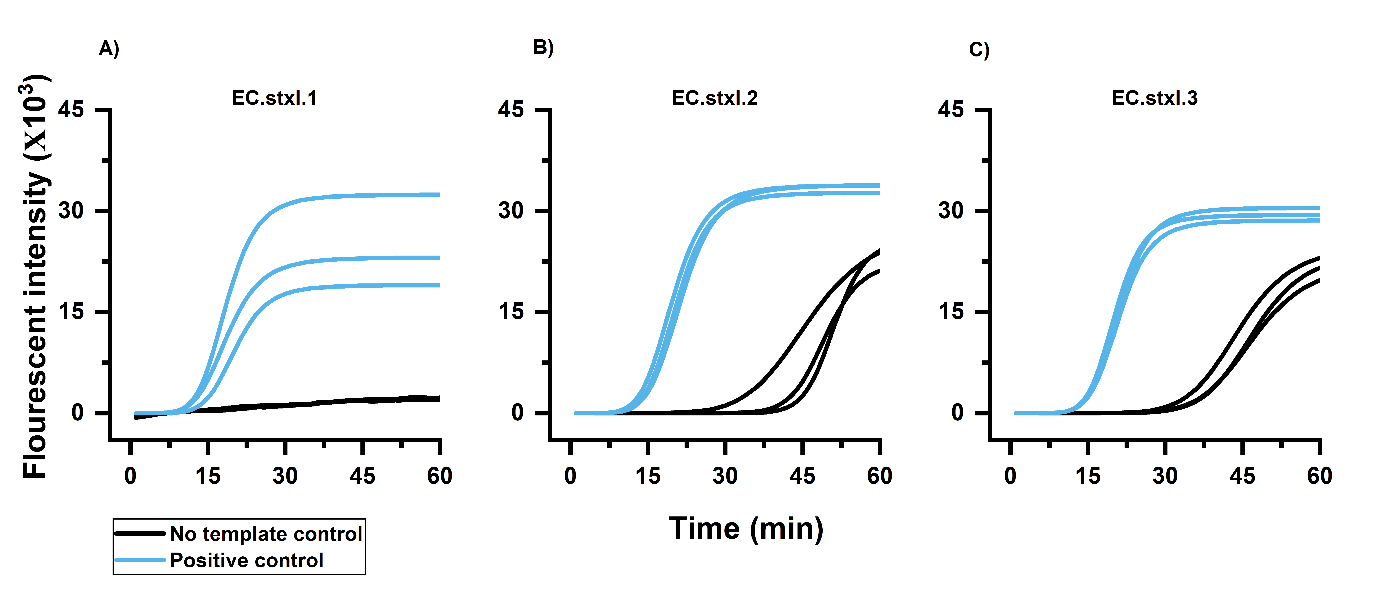

Figure S6: Fluorometric screening of primers for *stxI* gene in water using purified genomic DNA of *E. coli* O157:H7. The graphs are the LAMP amplification curves for varying primer sets in water. Blue lines indicate positive control where 5 µL of 0.2 ng/µL A) Primer set 1: EC.stxI.1, B) Primer set 2: EC.stxI.2, C) Primer set 3: EC.stxI.3. Black lines indicate non-template controls (NTC) where 5 µL of water was added. Three replicates of each condition were run per primer set.

On screening the three primer sets with genomic DNA of *E. coli* O157:H7, we observed that EC.stxI.1 primer set showed no amplifications in negatives, and the amplification start point in test reactions was within the desirable range of 15 minutes. The primer sets EC.stxI.2 and EC.stxI.3 showed quite early amplifications in negatives. We decided to proceed with EC.stxI.1. During our literature review, it was also observed that the sequence of these primers (Table S4) is nearly identical to the primer used by Wang *et al*. to detect *E. coli* O157:H7 in ground beef samples^1^.

Table S6: Sequence of primers used for PCR assays of *E. coli* O157:H7 *stxI* gene

| **Oligo sequence (5'- 3')** | **Oligo name** |
| --- | --- |
| GTGGCATTAATACTGAATTGTCATCA | Forward primer |
| GCGTAATCCCACGGACTCTTC | Reverse primer |
| TxRd-TGATGAGTTTCCTTCTATGTGTCCGGCAGAT-BHQ2 | Probe |

Table S7: Sequence of primers used for PCR assays of *Salmonella enterica* *invA* gene

| **Oligo sequence (5'- 3')** | **Oligo name** |
| --- | --- |
| AACGTGTTTCCGTGCGTAAT | Forward primer |
| TCCATCAAATTAGCGGAGGC | Reverse primer |
| FAM-TGGAAGCGCTCGCATTGTGG-BHQ | Probe |

2. **Supporting data for live-dead assay optimization of *E. coli* O157:H7**

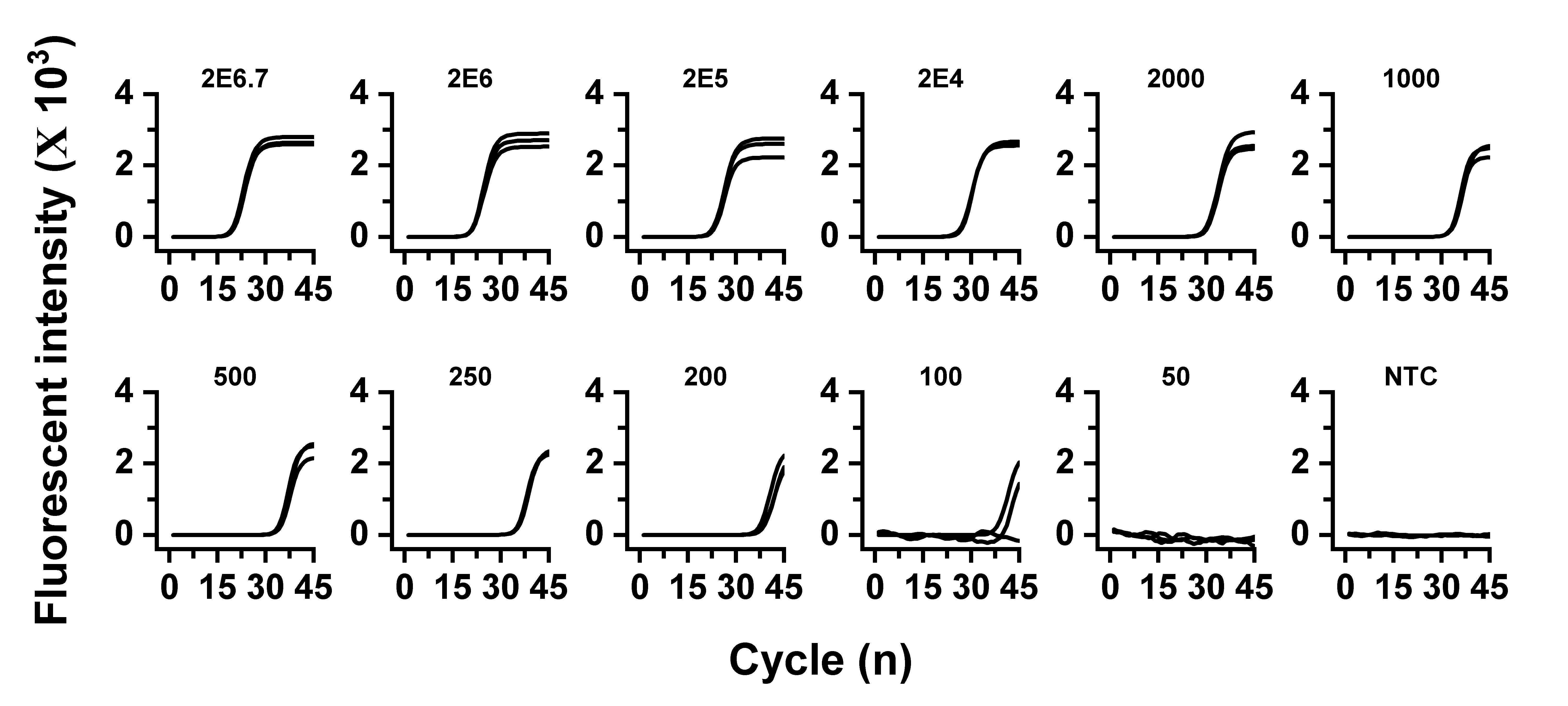

Figure S7: Fluorescent amplification data illustrating the limit of detection for PCR assays with whole dead *E. coli* O157:H7. The cell number in the reaction is labeled at the top of each graph. . Each sample was tested in triplicates.

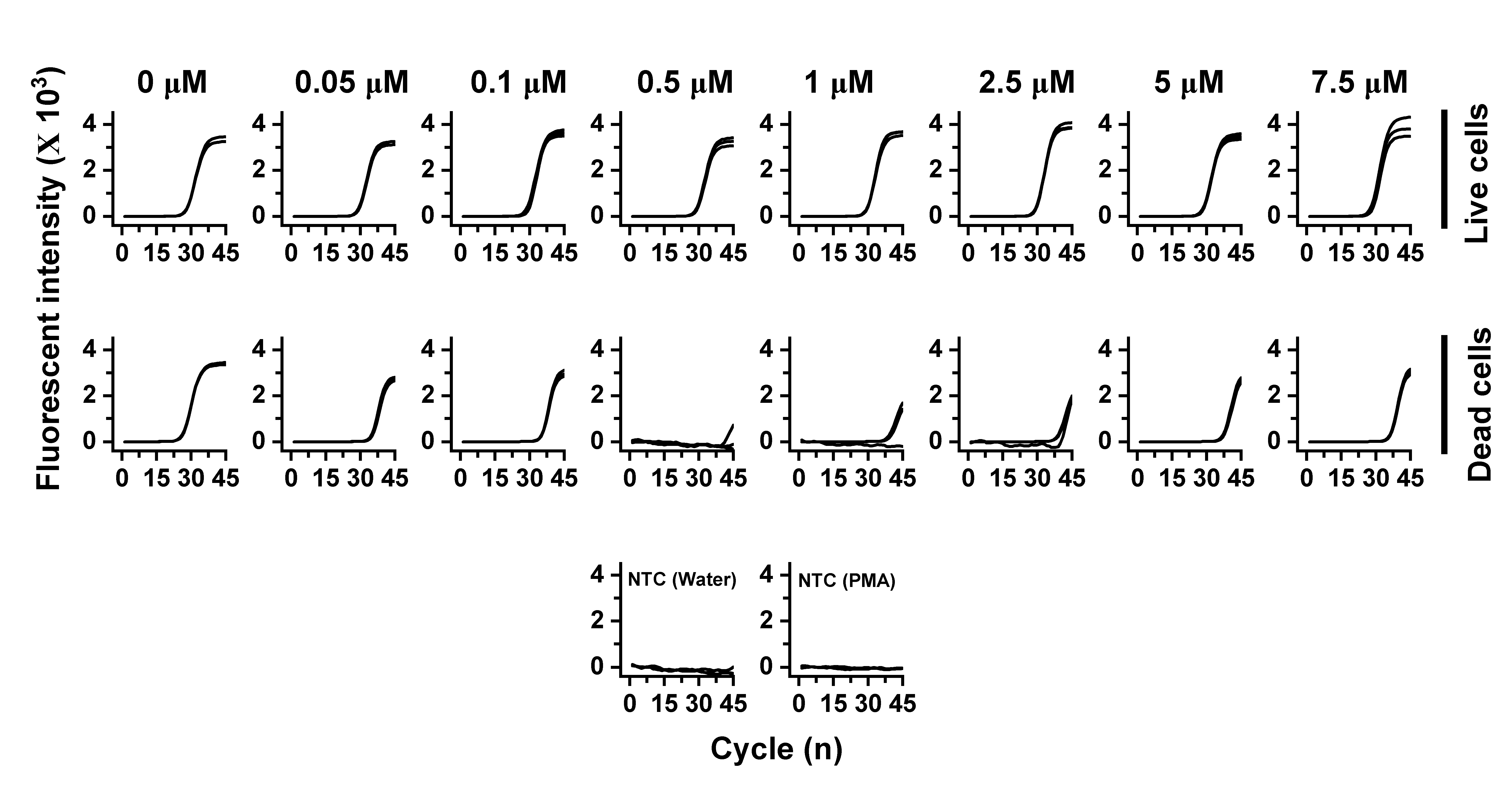

Figure S8: PCR-PMA results for different concentrations of PMA with a cell density of 2x10^6^ cells/mL. A) Fluorescent PCR amplification data for PMA-treated live cells (row 1 of graphs) and dead cells (row 2). The figure displays fluorescent data for both live and dead cells at each concentration of PMA, labeled at the top. Third row of graph presents negative controls with water and PMA solution as samples instead of bacterial cells. . Each sample was tested in triplicates. Each sample was tested in triplicates.

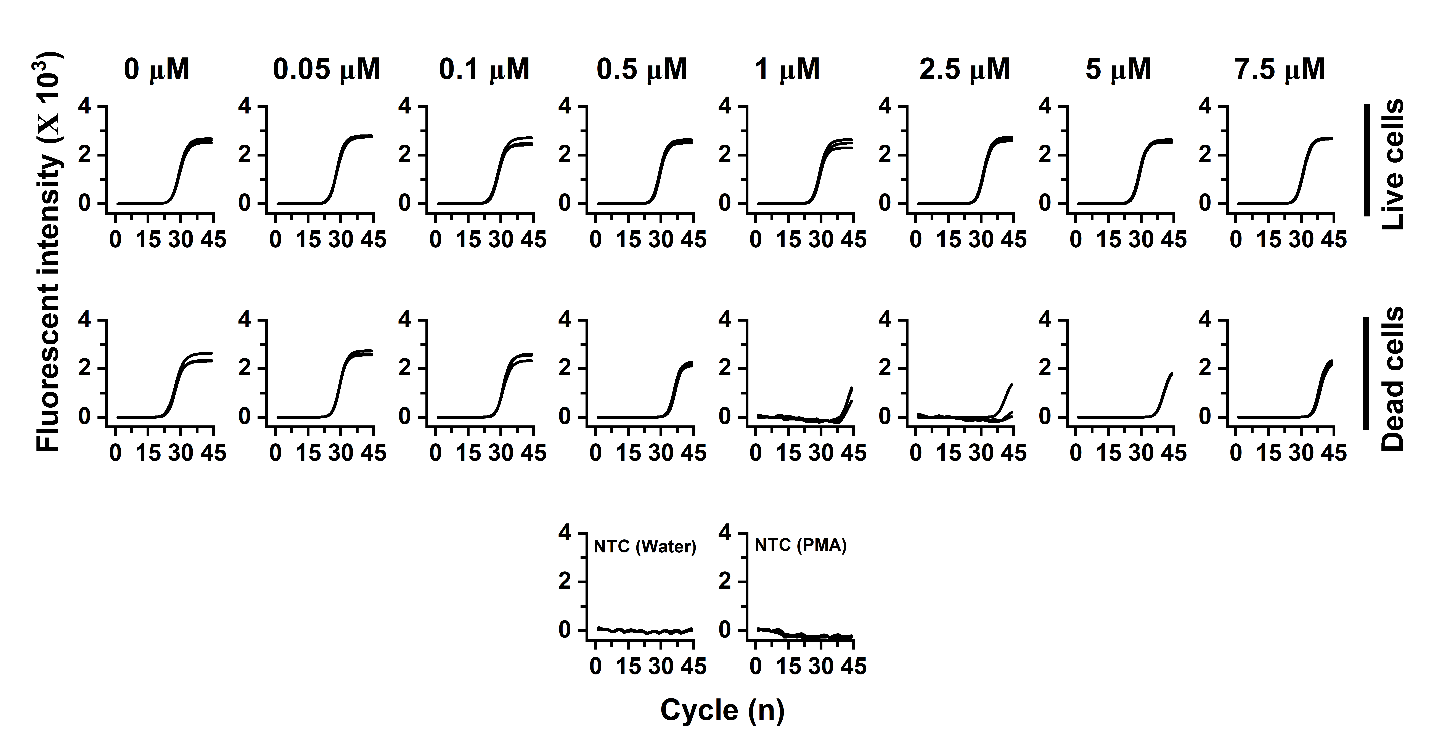

Figure S9: PCR-PMA results for different concentrations of PMA with a cell density of 2x10^7^ cells/mL A) Fluorescent PCR amplification data for PMA live cells (row 1 of graphs) and dead cells (row 2). The figure displays fluorescent data for both live and dead cells at each concentration of PMA, labeled at the top. Third row of graph presents negative controls with water and PMA solution as samples instead of bacterial cells. Each sample was tested in triplicates.

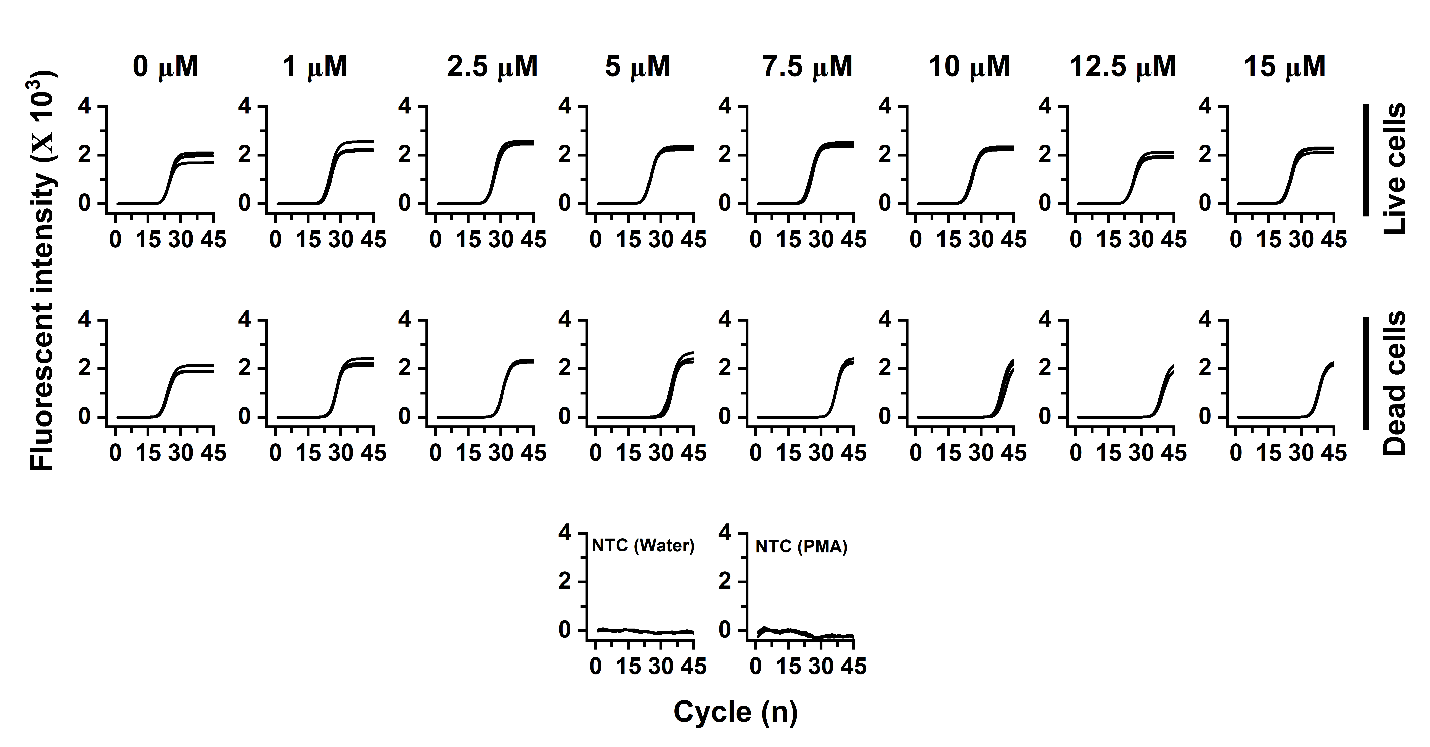

Figure S10: PCR-PMA results for different concentrations of PMA with a cell density of 2x10^8^ cells/mL A) Fluorescent PCR amplification data for PMA live cells (row 1 of graphs) and dead cells (row 2). The figure displays fluorescent data for both live and dead cells at each concentration of PMA, labeled at the top. Third row of graph presents negative controls with water and PMA solution as samples instead of bacterial cells. Each sample was tested in triplicates.

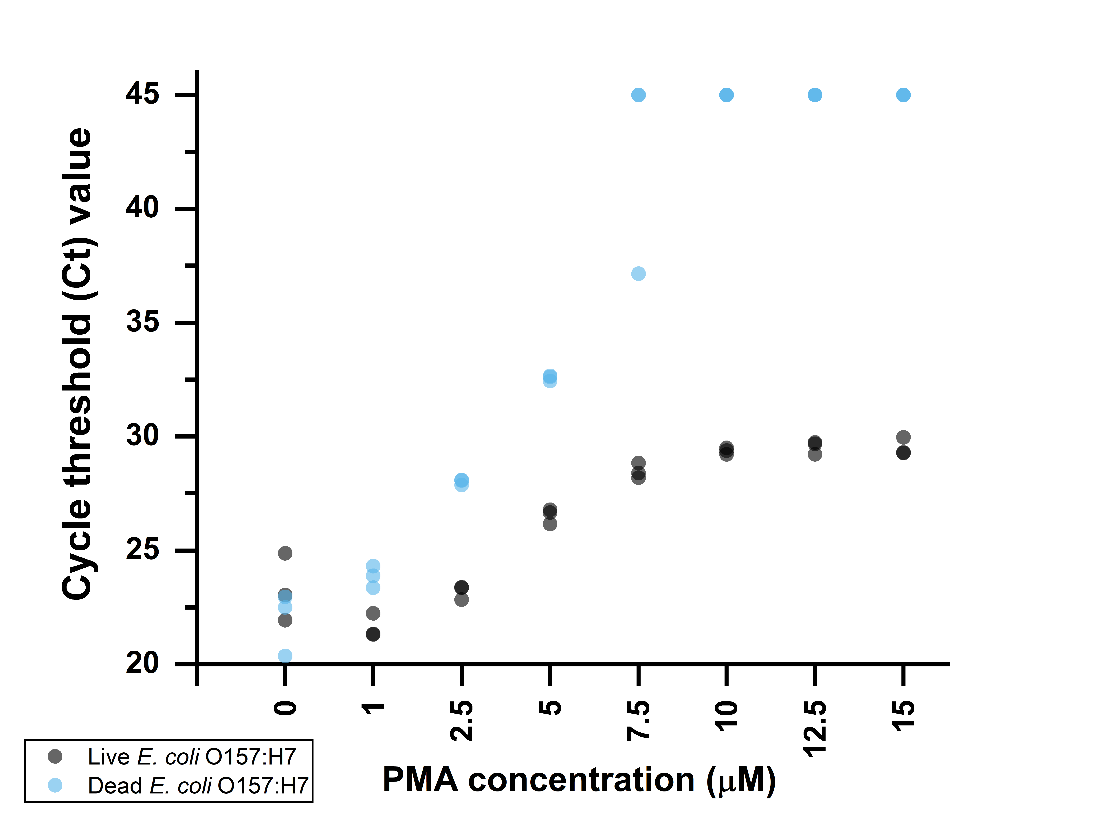

Figure S11: The plots show the cycle threshold (Ct) values obtained from qPCR analysis of PMA-treated cells and an additional precipitation step by centrifugation. Each sample was tested in triplicate. The x-axis represents the range of PMA concentrations used, while the y-axis represents the Ct values. This experiment is performed with 2x10^8^ cells/mL. In cases where no amplification was observed, the Ct value was assigned as the last cycle (45).

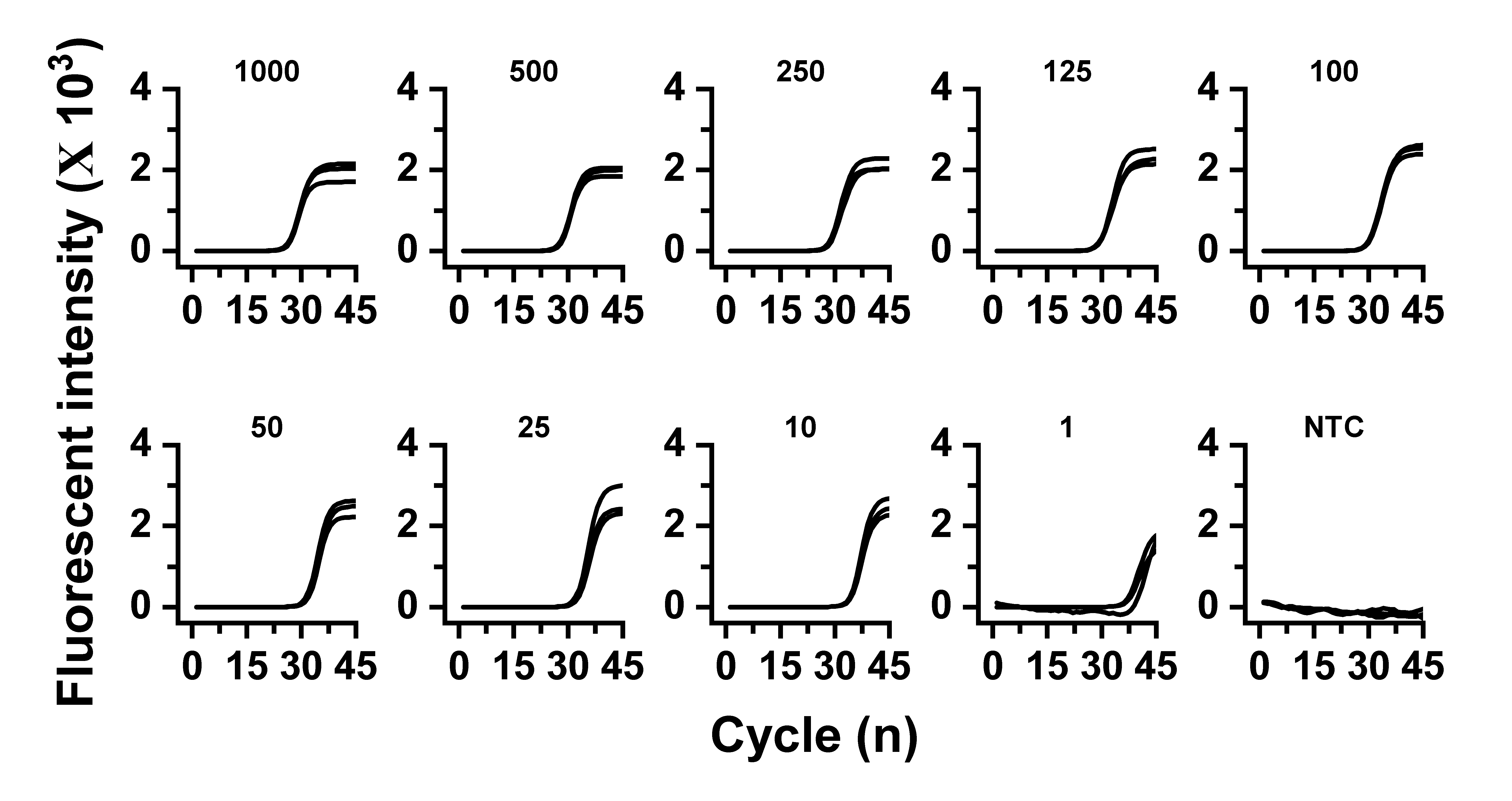

Figure S12: Fluorescent amplification data illustrating the limit of detection of PCR assays with genomic DNA of *E. coli* O157:H7. The DNA copy number in the reaction is labeled at the top of each graph. Each sample was tested in triplicates.

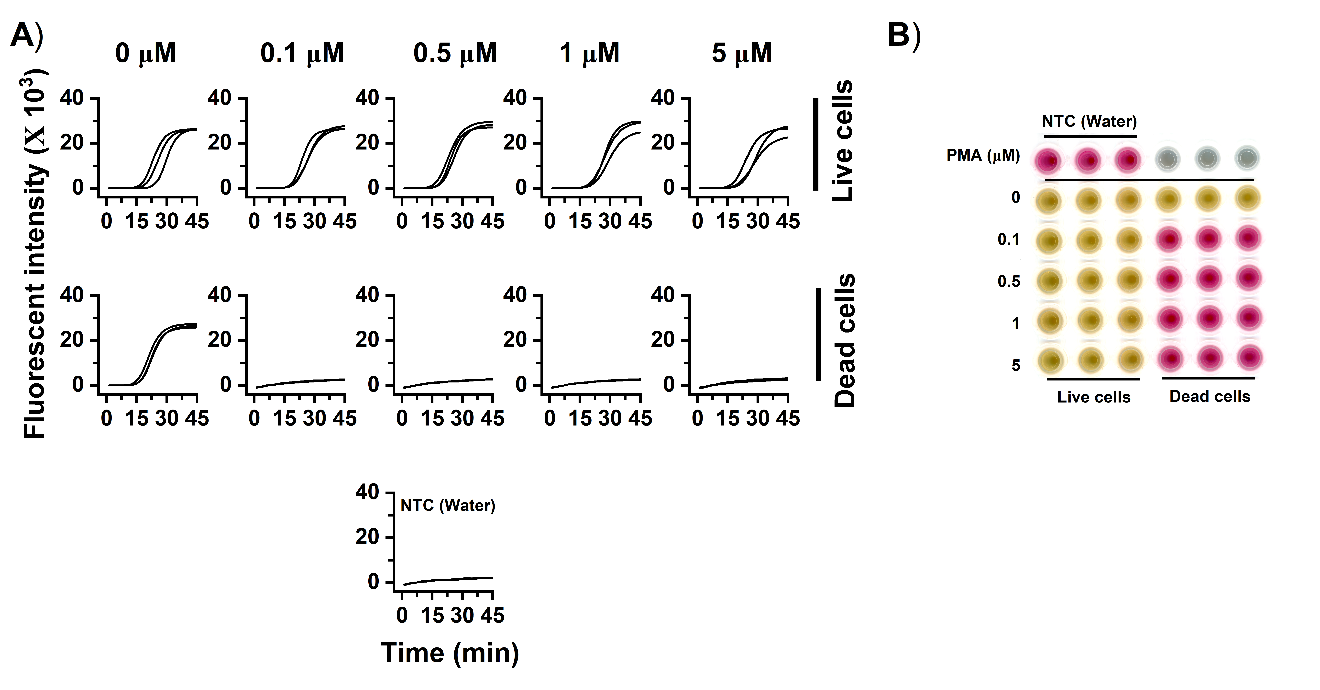

Figure S13: PMA-LAMP results for different concentrations of PMA with a cell density of 2x10^6^ cells/mL A) Fluorescent LAMP amplification data for PMA-treated live cells (row 1 of graphs) and dead cells (row 2). The figure shows fluorescent data for both live and dead cells at each concentration of PMA, labeled at the top. The third row of graph presents negative controls with water as samples instead of bacterial cells. Each sample was tested in triplicates. B) Scan of the microwell plate showing corresponding colorimetric data of LAMP using phenol red as an indicator. Change in color from red to yellow results from decreased pH due to the amplification reaction. Wells with no amplification appear red.

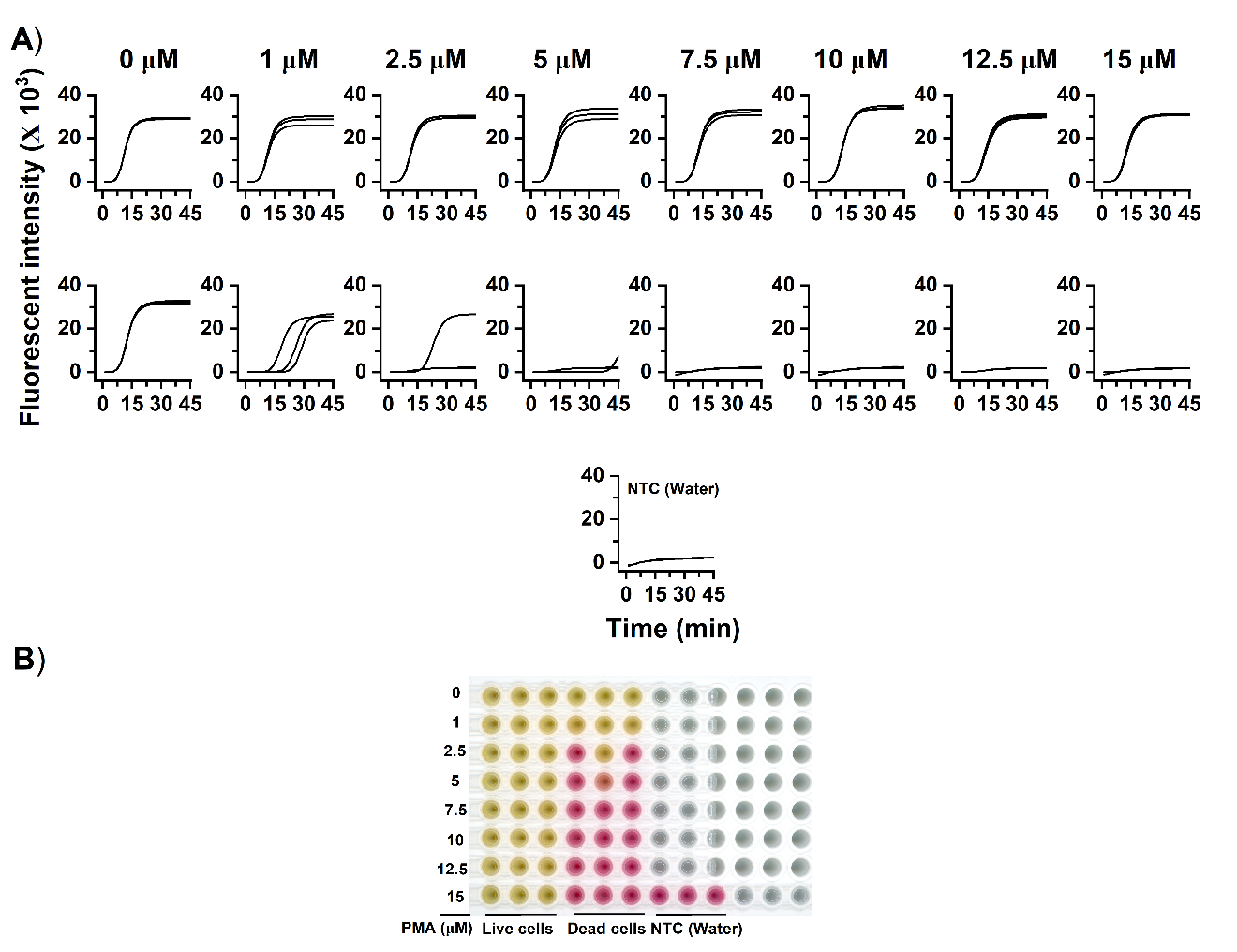

Figure S14: PMA-LAMP results for different concentrations of PMA with a cell density of 2x10^8^ cells/mL A) Fluorescent LAMP amplification data for PMA-treated live cells (row 1 of graphs) and dead cells (row 2). The figure shows fluorescent data for both live and dead cells at each concentration of PMA, labeled at the top. The third row of graph presents negative controls with water as samples instead of bacterial cells. Each sample was tested in triplicates. B) Scan of the microwell plate showing corresponding colorimetric data of LAMP using phenol red as an indicator. Change in color from red to yellow results from decreased pH due to the amplification reaction. Wells with no amplification appear red.

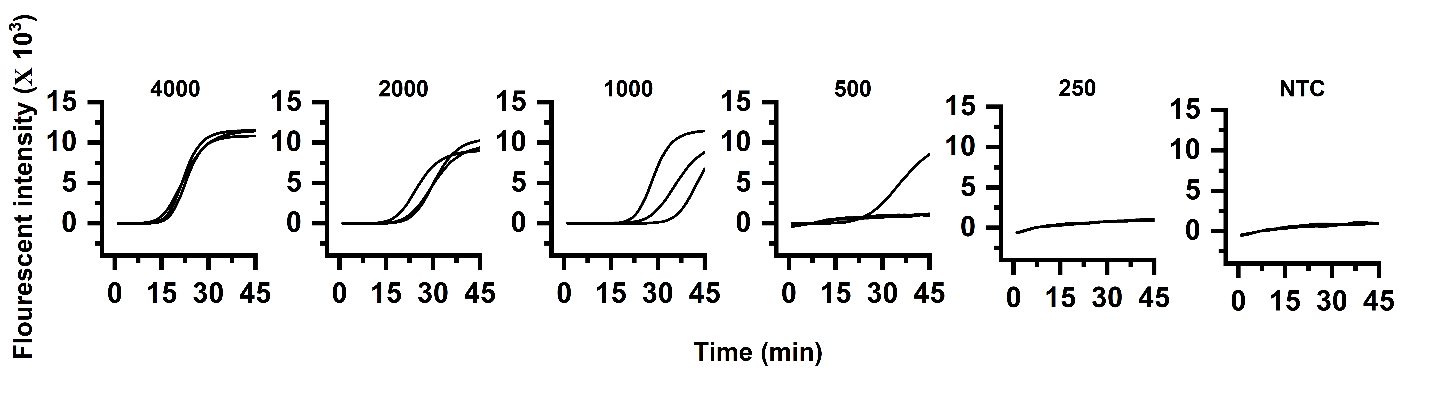

Figure S15: Fluorescent amplification data for limit of detection of LAMP assays with whole dead *E. coli* O157:H7. The cell number in the reaction is labeled at the top of each graph. Each sample was tested in triplicates.

3. **Supporting data for live-dead optimization of *Salmonella enterica***

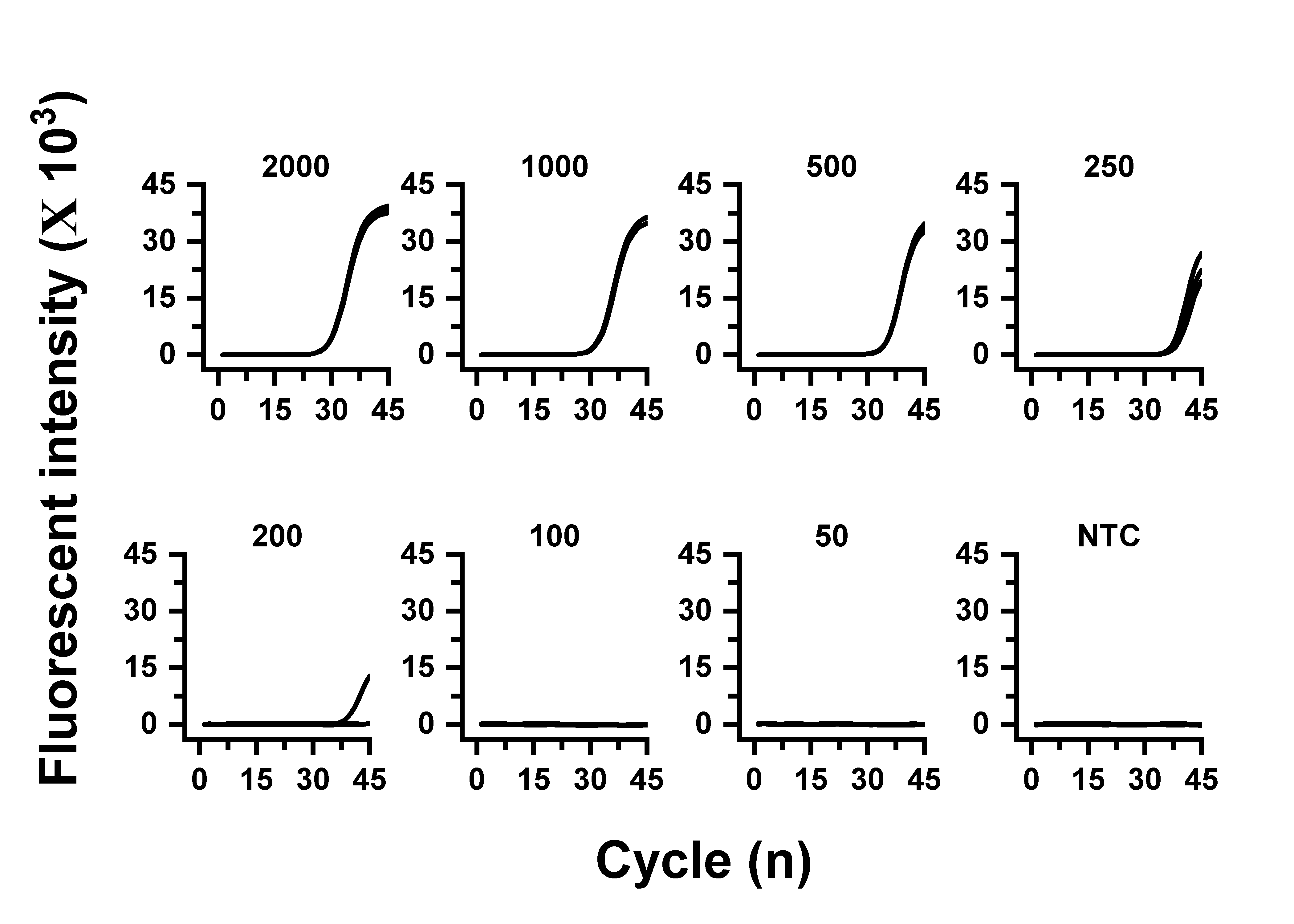

Figure S16: Fluorescent amplification data for the detection limit of LAMP assays with whole dead *Salmonella enterica*. The cell copy number in the reaction is labeled at the top of each graph. Each sample was tested in triplicates.

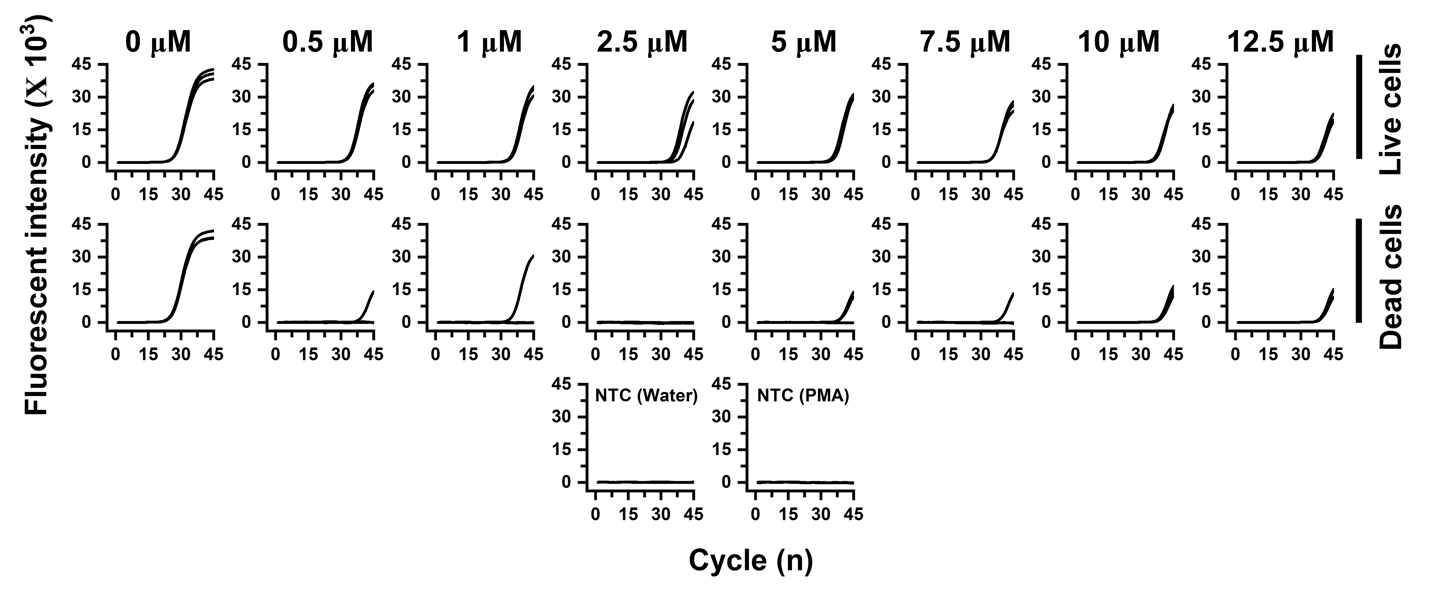

Figure S17: PCR-PMA results for different concentrations of PMA with a cell density of 2x10^6^ cells/Ml. A) Fluorescent PCR amplification data for PMA-treated live cells (row 1 of graphs) and dead cells (row 2). The figure displays fluorescent data for both live and dead cells at each concentration of PMA, labeled at the top. Each sample was tested in triplicates. Third row of graph presents negative controls with water and PMA solution as samples instead of bacterial cells.
